# CCT and Cullin1 regulate the TORC1 pathway to promote dendritic arborization in health and disease

**DOI:** 10.1101/2023.07.31.551324

**Authors:** Erin N. Lottes, Feyza H. Ciger, Shatabdi Bhattacharjee, Emily A. Timmins-Wilde, Benoit Tete, Tommy Tran, Jais Matta, Atit A. Patel, Daniel N. Cox

**Affiliations:** Neuroscience Institute, Georgia State University, Atlanta, GA 30303, United States

## Abstract

The development of cell-type-specific dendritic arbors is integral to the proper functioning of neurons within their circuit networks. In this study, we examine the regulatory relationship between the cytosolic chaperonin CCT, key insulin pathway genes, and an E3 ubiquitin ligase (Cullin1) in homeostatic dendritic development. CCT loss of function (LOF) results in dendritic hypotrophy in *Drosophila* Class IV (CIV) multidendritic larval sensory neurons, and CCT has recently been shown to fold components of the TOR (Target of Rapamycin) complex 1 (TORC1), *in vitro.* Through targeted genetic manipulations, we have confirmed that LOF of CCT and the TORC1 pathway reduces dendritic complexity, while overexpression of key TORC1 pathway genes increases dendritic complexity in CIV neurons. Both CCT and TORC1 LOF significantly reduce microtubule (MT) stability. CCT has been previously implicated in regulating proteinopathic aggregation, thus we examined CIV dendritic development in disease conditions as well. Expression of mutant Huntingtin leads to dendritic hypotrophy in a repeat-length-dependent manner, which can be rescued by TORC1 disinhibition via Cullin1 LOF. Together, our data suggest that Cullin1 and CCT influence dendritic arborization through regulation of TORC1 in both health and disease.

**SIGNIFICANCE:** The insulin pathway has become an increasingly attractive target for researchers interested in understanding the intersection of metabolism and brain health. We have found connections between the insulin pathway and cytosolic protein maintenance in the development of neuronal dendrites. These pathways converge on the dendritic cytoskeleton, particularly microtubules. Neurons expressing mutant Huntingtin also show defects in dendritic development and the underlying cytoskeleton, and we find that disinhibition of the insulin pathway can rescue dendritic hypotrophy in these neurons. This work advances our understanding of the molecular interactions between the insulin pathway and neuronal development in both health and Huntington’s Disease conditions.

## INTRODUCTION

It was once thought that the brain was isolated from the effects of starvation: at the start of the 20^th^ century, Edward H. Dewey, M.D. asserted, “The brain is not only a self-feeding organ when necessary, but it is also a self-charging dynamo, regaining its exhausted energies entirely through rest and sleep” (Dewey, 1900). Cognitive symptoms were repeatedly connected to diabetes (Miles and Root, 1922; Moheet et al., 2015), but it wasn’t until the discovery of neuronal insulin that the idea of the metabolically insulated brain was retired (Raizada, 1983; Weyhenmeyer and Fellows, 1983; Pardridge et al., 1985; Craft, 2009). Research linking neurodegenerative disorders to insulin resistance have since highlighted the necessity of understanding how the insulin pathway modulates the brain in health and disease (Schulingkamp et al., 2000; Wu et al., 2017; Kellar and Craft, 2020; Burillo et al., 2021). In this study, we examine putative connections among three cytosolic mechanisms – the insulin pathway, chaperone activity, and ubiquitin ligase activity – which each coordinate dendritic arborization through the regulation of the cytoskeleton.

In *Drosophila melanogaster*, the chaperonin CCT (<u>C</u>omplex <u>C</u>ontaining <u>T</u>ailless Complex Polypeptide 1 [TCP-1], also known as TRiC) is required for dendritic development of Class IV (CIV) multidendritic (md) sensory neurons of the larval peripheral nervous system (Das et al., 2017; Wang et al., 2020). CCT is a chaperonin – an ATP-dependent chaperone – and has the canonical chaperonin “barrel” shape, composed of two repeating rings of eight subunits each: CCT1-8 (**Fig 1A**) (Liou and Willison, 1997; Jin et al., 2019). Estimated to fold from 1-15% of cellular proteins, two of CCT’s most notable clients are cytoskeletal monomeric subunits actin and tubulin (Thulasiraman et al., 1999; Grantham et al., 2006; Brackley and Grantham, 2009; Willison, 2018; Gestaut et al., 2022).

**Figure 1:**
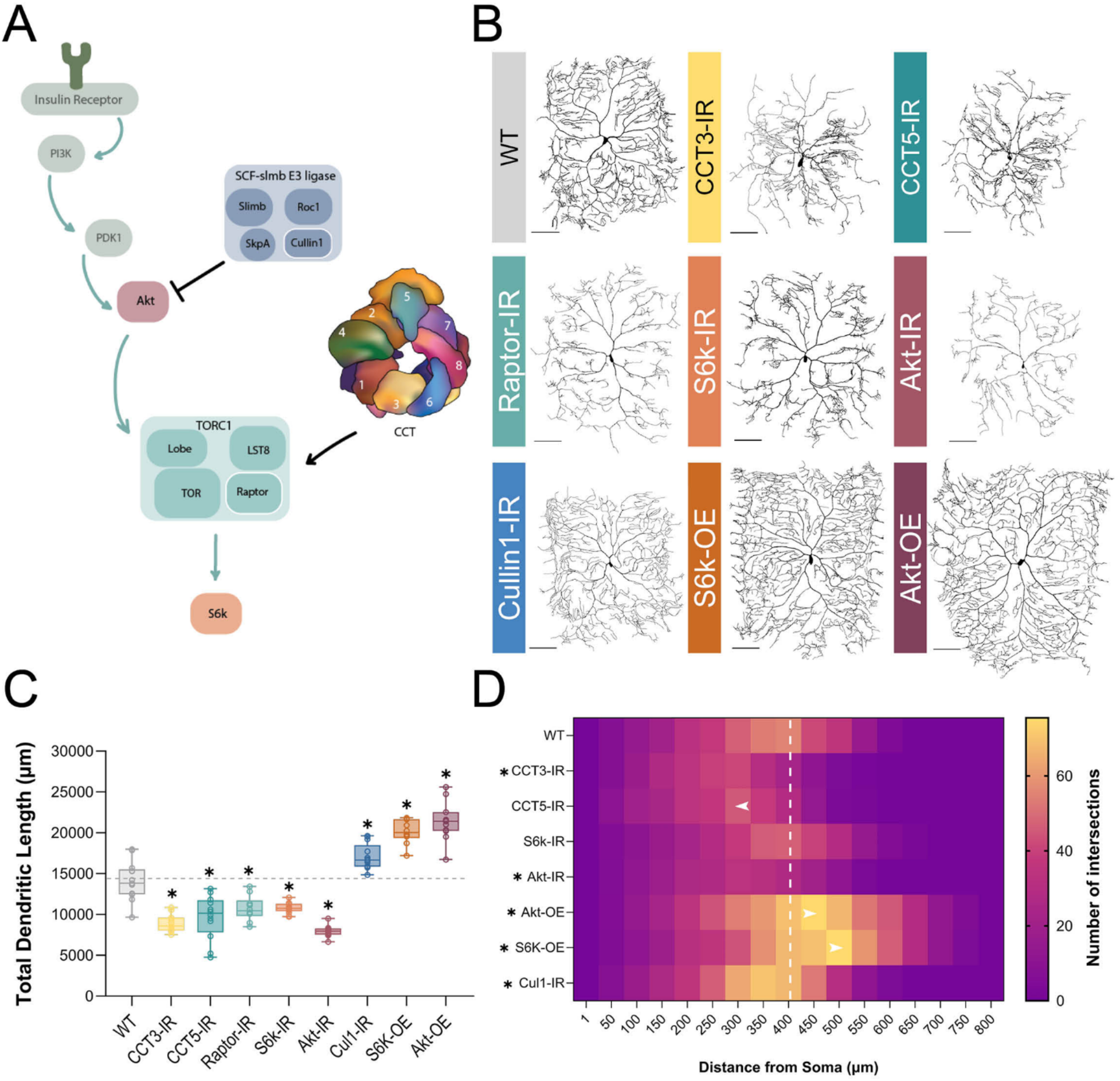
CCT and the TORC1 pathway promote dendritic arborization. (**A**) Schematic diagram of regulatory relationships between the insulin pathway, SCF complex, CCT, and TORC1 pathway with the insulin pathway indicated by teal arrows. TORC1 is negatively regulated by Cullin1 and positively regulated by CCT. The upstream insulin pathway in green is displayed for context but was not examined in this study. Individual components of the SCF and TORC1 complexes examined in this study are outlined in white. (**B**) Representative images of CIV neurons for key CCT and TORC1 pathway manipulations, with RNAi-mediated knockdown indicated with -IR and UAS-mediated overexpression with -OE. Scale bars = 100 μm (**C**) Total dendritic length of CCT and TORC1 pathway manipulations shown in comparison to a WT control, * indicates a significant change from control (p < 0.05). (**D**) Number of Sholl intersections mapped by color at increasing radial distances from soma (μm). Significant changes in Sholl maximum intersections and radius are indicated by an asterisk. Arrows indicate genotypes where the radius of maximum intersections has shifted significantly from WT. The white dashed line is to reference the radius of maximum intersections for WT neurons. In all panels * = p < 0.05, see **Supplementary Table S2** for detailed statistics.

CCT physically interacts with proteins in the TORC1 pathway and folds Raptor, the regulatory component of TORC1 (Cuéllar et al., 2019; Kim and Choi, 2019). TORC1 is a part of the insulin pathway, operating downstream of Phosphatidylinositol-3 kinase (PI3K) and Akt kinase (Fig 1A), and has been long known to control cell size (Fingar et al., 2002; Kumar et al., 2005) as well as dendritic development in many organisms (Jaworski et al., 2005; Kumar et al., 2005; Lee and Chung, 2007; Urbanska et al., 2012; Thomanetz et al., 2013; Shimono et al., 2014; Skalecka et al., 2016; Kosillo et al., 2019, 2022).

In this study, we investigate the regulatory roles of CCT, TORC1, and the SCF complex in both conditions of homeostatic development and proteinopathic disease states. Cullin1 is the scaffolding component of the <u>S</u>kp, <u>C</u>ullin, <u>F</u>-box (SCF) E3-ubiquitin ligase – previously shown to regulate TORC1 in dendritic pruning in *Drosophila* (Wong et al., 2013). In homeostasis, via *in vivo* analyses of CIV neurons, we establish that dendritic arborization is mediated by a chaperonin (CCT) and E3 ubiquitin ligase component (Cullin1), both of which partially mediate dendritic complexity through regulation of TORC1. TORC1 inhibition leads to dendritic hypotrophy whereas TORC1 activation leads to dendritic hypertrophy.

CCT and TORC1 have also been examined as endogenous mediators of the cellular effects of Huntington’s Disease (HD), a neurodegenerative disease caused by a polyglutamine expansion mutation. Targeted manipulations of CCT and TORC1 have been found to reduce aggregates and enhance cell viability in multiple model systems of HD (Sontag et al., 2013; Lee et al., 2015; Noormohammadi et al., 2016; Shen et al., 2016). TORC1 inhibition, via application of rapamycin and similar drugs, has been shown to be neuroprotective in cell culture models of HD, as well as in both *Drosophila* and zebrafish photoreceptors with mutant Huntingtin (Ravikumar et al., 2004; Berger et al., 2006; King et al., 2008; Williams et al., 2008). However, the role of TORC1 and CCT in regulating dendritic development in HD conditions was previously unexplored. In CIV neurons, although high repeat mHTT expression results in dendritic hypotrophy and loss of underlying microtubule signal like that of TORC1 and CCT LOF, we do not find evidence that mHTT disrupts the TORC1-CCT dendritic arborization pathway in HD conditions.

## METHODS

### *Drosophila* husbandry and stocks

The *Drosophila melanogaster* stocks used in this study were reared on a standard recipe of cornmeal, molasses, and agar media and maintained at 25°C. Genetic crosses for live imaging and immunohistochemistry were reared at 29°C. In all experiments, larvae were randomized for sex. A complete list of stocks and genetic lines for this study is provided in **Supplementary Table S1**.

### Immunohistochemical analysis

Larval dissection, mounting, and staining were performed as previously described (Grueber et al., 2002; Sulkowski et al., 2011). Larvae stained for CCT, acetylated α-tubulin, and Raptor were dissected and muscles removed as previously described (Tenenbaum and Gavis, 2016), and fixed samples were imaged with a Zeiss LSM780 Confocal microscope under 63X magnification using an oil-immersion objective. Primary antibodies used include: chicken anti-GFP (1:1000 dilution, Aves Labs), rabbit anti-phosphorylated S6k (1:300 dilution, Cell Signaling Technology), rabbit anti-Raptor (1:200 dilution, Cell Signaling Technology), mouse anti-acetylated α-tubulin (1:100 dilution, Santa Cruz Biotechnology), mouse anti-Futsch (1:100 dilution, Developmental Studies Hybridoma Bank), rabbit anti-Huntingtin (1:200 dilution, Cell Signaling Technology), rabbit anti-HA (1:500 dilution, Cell Signaling Technology), rat anti-CCT1 (1:200 dilution, Origene), mouse anti-CCT5 (1:200 dilution, GeneTex), mouse anti-S6k (1:200 dilution, Proteintech), rabbit anti-phosphorylated Akt (1:200 dilution, Cell Signaling Technology), rabbit anti-Cullin1 (1:200 dilution, Invitrogen), mouse anti-β-tubulin IIA (1:500 dilution, Novus Biologicals). Secondary antibodies used include: donkey anti-chicken 488 (1:2000 dilution, Jackson Immunoresearch), donkey anti-mouse 555 (1:200 dilution, Life Technologies), donkey anti-mouse 568 (1:200 dilution, Life Technologies), donkey anti-mouse 647 (1:200 dilution, Life Technologies), donkey anti-rabbit 568 (1:200 dilution, Life Technologies), donkey anti-rabbit 647 (1:200 dilution, Life Technologies), donkey anti-rat Cy3 (1:200 dilution, Jackson Immunoresearch).

### Live confocal imaging, neural reconstructions, and morphometric analyses

Live imaging was performed using the Zeiss LSM780 Confocal as previously described (Iyer et al., 2013a, 2013b). Multiple gene-specific RNAi lines were examined for each genotype and validated using IHC and mutant analysis when possible. MARCM analysis was performed as previously described (Sulkowski et al., 2011; Iyer et al., 2013b). To generate CIV neuron MARCM clones *GAL^5-40^UAS-Venus:pm SOP-FLP#42;tubP-GAL80FRT40A* (2L MARCM) flies were crossed to *CCT4^KG09280^,FRT40A* mutant flies. Maximum intensity projections of dendritic z-stacks were processed and neurons reconstructed as previously described (Clark et al., 2018). Quantitative morphological data (including total dendritic length and Sholl analysis) were compiled using the Simple Neurite Tracer (SNT) plugin for FIJI (Ferreira et al., 2014; Arshadi et al., 2021). Batch processing was completed using a custom FIJI macro and Rstudio script created by Dr. Atit A. Patel (Cox Lab) and the resulting data was exported to Excel (Microsoft).

### Live multichannel neural reconstructions

Multichannel cytoskeletal reconstructions and related quantitative analyses were performed using the method described in (Nanda et al., 2021) and implemented in (Bhattacharjee et al., 2022) for CIV cytoskeletal analysis. In brief, one primary branch and all connected distal branches in the same posterior quadrant were reconstructed for each neuron using Neutube (Feng et al., 2015), then microtubule (MT) fluorescence was measured at distinct points along the dendritic arbor, averaged in 20 or 40 µm bins, and then normalized to 1 for comparison to controls. Displayed *Jupiter::mCherry* fluorescence is shown as a ratio of normalized fluorescence over the total path length within each bin, and can be understood as the average normalized fluorescence along a single branch.

### Co-localization analysis

Colocalization was performed on samples prepared via immunohistochemistry and imaged at 63X resolution under an oil immersion lens. A theoretical point spread function (PSF) was created for each measured wavelength using FIJI’s PSF Generator and the Born and Wolf 3D optical model and images were deconvolved with the FIJI DeconvolutionLab2 plugin using the Richardson-Lucy algorithm (Sage et al., 2017). Cytosolic compartments of interest were traced and regions of interest created using FIJI, then analyzed with BIOP JacoP to produce Pearson’s Correlation and Mander’s Coefficients (Bolte and Cordelières, 2006). Automatic thresholding based on percentile was used for Mander’s Coefficients for CCT and wild-type (WT) HTT signal. Manual thresholding for mHTT120-HA signal was based on the maximum intensity of off-target rabbit anti-HA signal in matched controls.

### Temperature-induced mHTT expression

mHTT overexpression lines were made by crossing *ppkEGFP,tsGAL80;ppkGAL4* with *UAS*-*mHTTQ120-HA* overexpression lines. *tsGAL80* binds *GAL4* at temperatures <29°C and prevents transcription of *UAS*-driven mHTT misexpression (McGuire et al., 2004). However, at temperatures ≥29°C, *tsGAL80* cannot bind *GAL4* which allows *mHTTQ120-HA* overexpression. *ppkEGFP,tsGAL80;ppkGAL4* was crossed to *Oregon R* (WT) and *UAS-mHTTQ120-HA* at RT (25°C) and allowed to lay eggs for three hours. Larvae were raised in two conditions: 96 hours at 25C (*GAL4* “off”) then stained for HTT (WT HTT), or 48 hours at 25°C followed by 48 hours at 30°C (*GAL4* “on”) and stained for HA (HTT120). Larvae were dissected 96 hours after egg laying (AEL) as described above. Dissected larvae were stained for HTT and CCT1 or HA and CCT1 and prepared and imaged as described above for all other immunohistochemical analyses.

### mHTT aggregate inclusion body analysis

Inclusion body aggregates of mHTT were manually quantified using Zen Blue Lite software in neurons expressing *mHTTQ96-Cerulean* imaged live at 20X magnification. Neuron labels were coded for analysis to ensure blind conditions. Inclusion bodies were outlined using the “Draw Spline Contour” tool and average area compared across genotypes.

### Experimental design and statistical analyses

Statistical analyses were performed using GraphPad Prism 10. Error bars in the figures represent standard error of the mean (SEM). All data were tested for normality using the Shapiro-Wilk normality test. Statistical tests performed include: unpaired *t*-test; one-way ANOVA with Dunnett’s, Šídák’s, or Tukey’s multiple comparison test (multiple comparison tests chosen based on Prism 10 recommendations); two-way ANOVA with Tukey’s multiple comparison test; Mann-Whitney *U* test; Kruskal Wallis test using Dunn’s multiple comparison test. Data points lying greater than two standard deviations above or below the mean were removed. A single asterisk is used in all graphs to denote significance (p ≤ 0.05), and detailed statistical results are available in **Supplementary Table S2**.

## RESULTS

### CCT LOF and of disruption of TORC1 pathway genes results in dendritic hypotrophy

CCT is required for complex dendritic arbor formation in *Drosophila melanogaster* CIV md neurons (Das et al., 2017; Wang et al., 2020), as we have independently confirmed in this study via RNAi and MARCM (**Fig 1B-D, S1A-D**). LOF of *CCT3* and *CCT5* both result in significant dendritic hypotrophy and will be used throughout this study to disrupt the CCT complex. LOF of single CCT subunits has previously been found to reduce expression of other CCT subunits (Freund et al., 2014; Chen et al., 2018; Kim and Choi, 2019), and we have independently found via immunohistochemistry (IHC) that *CCT4* LOF results in significant reductions in CCT5 expression (**Fig S1E**). Additionally, simultaneous RNAi knockdown of *CCT4* and *CCT5* results in significantly lower CCT5 expression than *CCT4-IR* or *CCT5-IR* knockdowns alone (**Fig S1E**). Developmental time course analyses reveal that CIV dendritic hypotrophy with *CCT3* or *CCT5* knockdowns first manifests at 72 hours after egg laying (AEL) – as indicated by reductions in total dendritic length (TDL) that plateau later in larval development (**Fig S1F**).

Though many clients of CCT have been identified, few have been examined alongside CCT in dendritic development. There is evidence that CCT folds components of TORC1 *in vitro* and co-operates with the insulin pathway to regulate organ size in *Drosophila* (Cuéllar et al., 2019; Kim and Choi, 2019). Using RNAi, we knocked down key insulin pathway effector genes: *Akt, Raptor,* and *S6k* – an activator, a component, and a downstream effector of TORC1, respectively (**Fig 1A**). Efficacy of various RNAi knockdowns was confirmed by quantifying fluorescence of each protein of interest in the wild-type (WT) control and RNAi knockdown conditions. *Raptor* RNAi results in significant reductions in Raptor fluorescence relative to control (**Fig S2A**). *S6k* RNAi (*S6k-IR*) likewise results in significant decreases in S6k (**Fig S2B**), and phosphorylated S6k (P-S6k) expression, the active form of S6k (**Fig 2A**). *Akt-IR* also results in a significant decrease of phosphorylated Akt expression (**Fig S2C**). Akt, Raptor, and S6k LOF all result in significant dendritic hypotrophy, as measured by total dendritic length (TDL) (**Fig 1B-C**). Furthermore, the maximum number of Sholl intersections is significantly decreased in *Akt* and *CCT3* knockdown conditions (**Fig 1D**), indicative of decreased branch complexity; additionally, *CCT5* knockdown leads to a significant shift in maximum radius toward the soma (**Fig 1D**). Collectively, these data demonstrate both CCT and the TORC1 pathway are required for CIV dendritic development.

**Figure 2:**
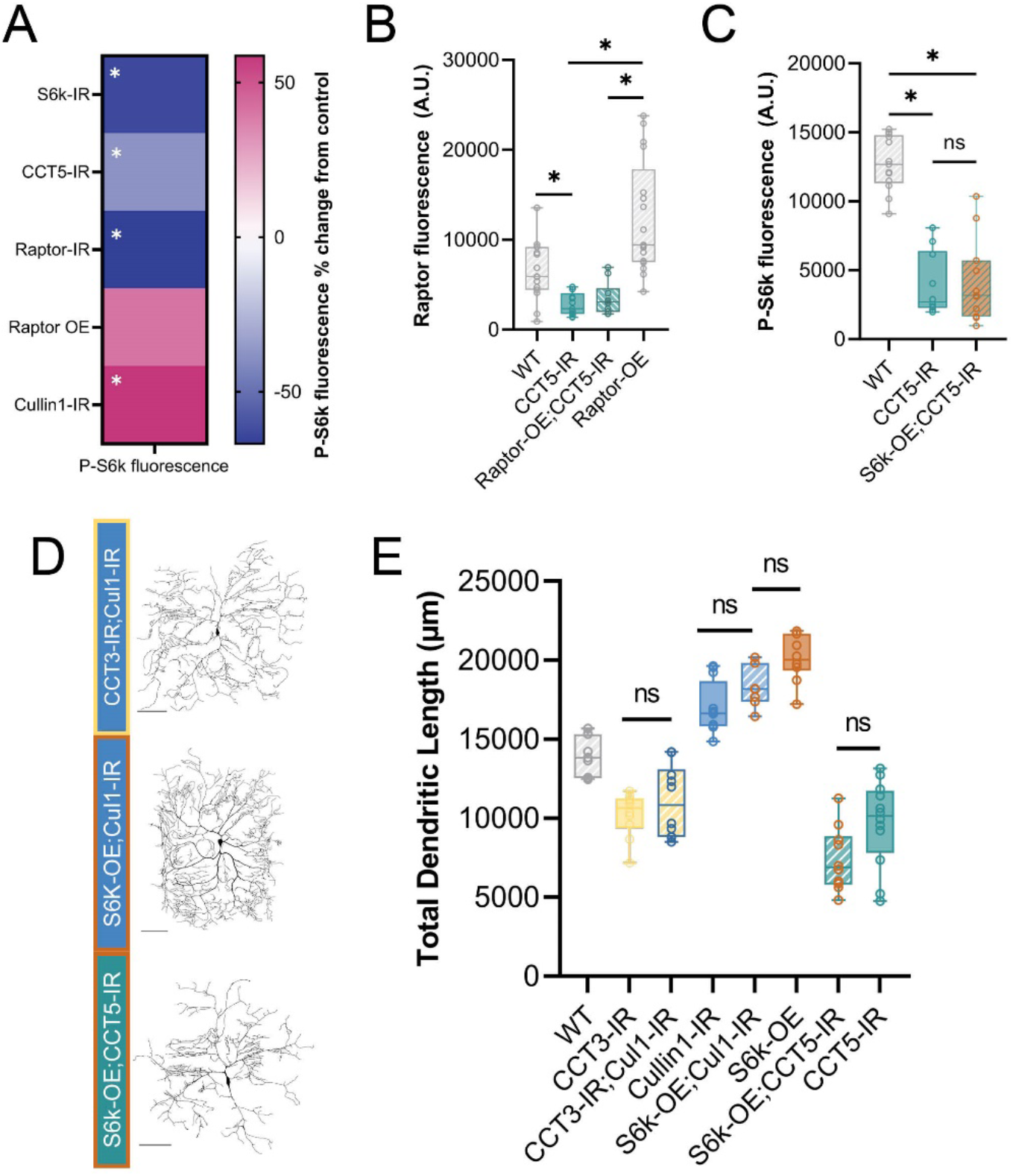
CCT and Cullin1 regulate the TORC1 pathway *in vivo*. (**A**) Heat map showing the percent change in P-S6k fluorescence for each genetic manipulation as compared to its proper control. (**B**) Raptor fluorescence is significantly decreased in *CCT5* LOF conditions and is not rescued by overexpression of Raptor. (**C**) P-S6k fluorescence levels are significantly decreased in *CCT5* LOF and are not rescued by overexpression of S6k. (**D**) Representative images of combined TORC1 genetic manipulations. Scale bars = 100 µm. (**E**) Total dendritic length in microns for WT and combined TORC1 genetic manipulations. In all panels * = p < 0.05, see **Supplementary Table S2** for detailed statistics.

### TORC1 hyperactivation results in dendritic hypertrophy

There is evidence in multiple model organisms that TORC1 over-activation can result in increases in dendritic complexity (Diaz-Ruiz et al., 2009; Kosillo et al., 2019; Kanaoka et al., 2023). In CIV neurons, overexpression (OE) of key components of the TORC1 pathway result in increased complexity of dendritic arbors (**Fig 1B-D**). *S6k OE* and *Akt OE* both result in significantly increased TDL, a significantly higher Sholl maximum, and a significantly, distally shifted Sholl maximum radius relative to controls (**Fig 1B-D**). In contrast, overexpression of individual subunits of CCT does not produce any significant change in dendritic TDL (**Fig S1G**), consistent with prior studies (Das et al., 2017). Overexpression of single CCT subunits has been found to be insufficient to increase levels of other CCT subunits in the complex (Tam et al., 2006; Noormohammadi et al., 2016).

Cullin1, a scaffolding component of the SCF complex, has previously been shown to regulate TORC1 activity in CIV dendrite pruning (Wong et al., 2013) through inhibition of Akt (**Fig 1A**). Another component of SCF, SkpA, was also previously reported to produce dendritic hypertrophy under LOF conditions (Das et al., 2017). We find that as in *SkpA* LOF and *Akt* OE, *Cullin1* LOF results in significantly increased TDL in CIV neurons (**Fig 1B-C**). Efficacy of *Cullin1* RNAi was confirmed via IHC as *Cullin1* LOF leads to a significant reduction in Cullin1 fluorescence in CIV neurons (**Fig S2D**).

To validate activation or inhibition of TORC1 through genetic manipulations of pathway components and cytosolic interactors, we stained for the downstream product of TORC1: phosphorylated S6k (P-S6k) (**Fig 1A, 2A**). LOF of *Raptor* and *CCT5* significantly decrease P-S6k levels, confirming disruption of TORC1 (**Fig 2A**). *Cullin1 LOF* significantly increases levels of P-S6k (**Fig 2A**) indicating that knockdown of *Cullin1* disinhibits TORC1 to phosphorylate S6k.

### CCT regulates Raptor levels *in vivo*

CCT was recently demonstrated to fold Raptor, the regulatory component of TORC1 (Cuéllar et al., 2019). We confirm that this regulatory relationship is conserved in *Drosophila melanogaster* larval sensory neurons. First, we find that *CCT5* knockdown significantly decreases levels of Raptor in CIV neurons from both WT and Raptor overexpression (**Fig 2B, S2A**). Though overexpression of Raptor via *UAS-Raptor-HA* significantly increases Raptor fluorescence over WT levels (**Fig S2A**), it is insufficient to increase Raptor fluorescence levels significantly in a *CCT5* LOF background, as measured via IHC (**Fig 2B**).

To further confirm the requirement of CCT to sustain the TORC1 pathway, we overexpress *S6k* in the *CCT5-IR* background (**Fig 2 C-E**). We hypothesized that if CCT were required for TORC1-mediated phosphorylation of S6k, then overexpression of S6k would not be sufficient to rescue either P-S6k levels or dendritic arborization. Indeed, overexpression of S6k could not return levels of P-S6k fluorescence to WT conditions in a *CCT5-IR* background (**Fig 2C**). Likewise, overexpression of S6k in *CCT5-IR* neurons was unable to rescue *CCT5* LOF-induced dendritic hypotrophy (**Fig 2D-E**) revealing CCT is necessary for *S6k* OE-induced dendritic hypertrophy. These data indicate that CCT is required for Raptor expression and subsequent S6k phosphorylation through TORC1.

### Cullin1 regulates dendritic arborization through TORC1

*Cullin1* knockdown significantly increases levels of P-S6k fluorescence as measured through IHC (**Fig. 2A**). *Cullin1* LOF and *S6k* overexpression both lead to dendritic hypertrophy (**Fig 1 B-D**). However, combining *Cullin1* knockdown and *S6k* overexpression in the same neurons does not further increase dendritic complexity, and there is no significant difference in TDL between the individual manipulations and the combined phenotype (**Fig 2D-E**).

We hypothesized that if *Cullin1* LOF were causing dendritic hypertrophy through regulation of TORC1, then it would be unable to recover any of the lost complexity in CCT LOF neurons, as we have established CCT is essential for TORC1 activity (**Fig 2B-C**). *Cullin1* knockdown does not significantly change TDL in a *CCT3* LOF background compared to *CCT3* knockdown alone (**Fig 2E**), indicating that CCT function is required for the observed hypertrophy in *Cullin1* knockdown neurons (**Fig 2E**).

### TORC1 pathway disruption results in loss of stable microtubules

CCT has been demonstrated to directly fold both α- and β-tubulin (Llorca, 2000; Gestaut et al., 2022) and regulate stable microtubule (MT) levels in CIV dendrites (Das et al., 2017; Wang et al., 2020). We have independently confirmed that CCT LOF leads to significant reductions in underlying levels of stable MTs through several measures (**Figs. 3A-C, S2E**). Since inhibition of the TORC1 pathway caused significant dendritic hypotrophy, and TORC1 has been previously linked to cytoskeletal phenotypes (Swiech et al., 2011), we predicted that there would be underlying cytoskeletal changes accompanying the loss of complexity.

**Figure 3:**
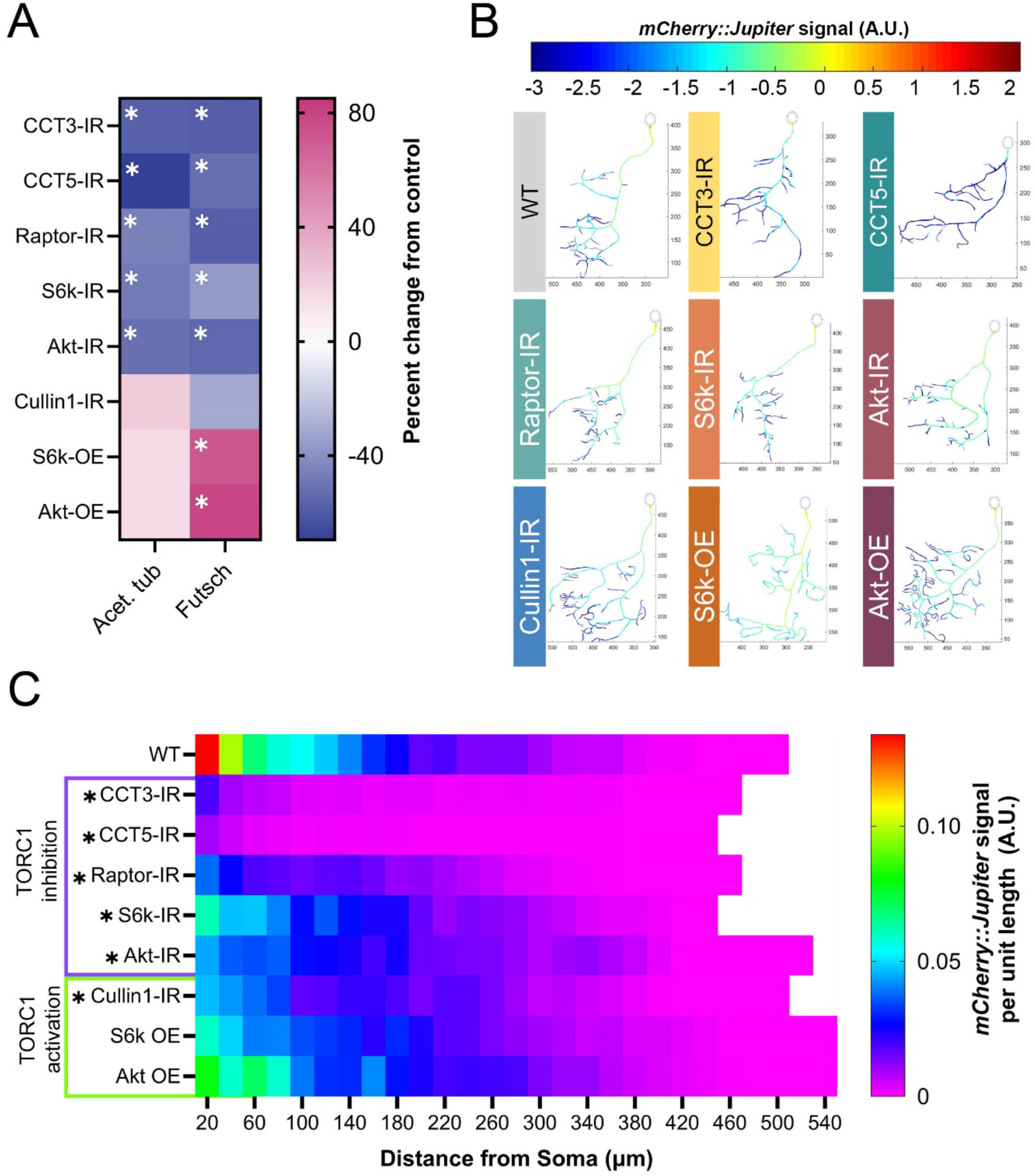
TORC1 pathway manipulations alter underlying stable MT signal. (**A**) Heat map showing percent change from control in acetylated α-tubulin and Futsch levels for each genetic manipulation. Each experimental condition was compared to WT control and appropriate statistical comparisons were performed (detailed in **Supplementary Table S2**). (**B**) Representative reconstructions of branches from WT and TORC1 genetic manipulations – normalized *mCherry::Jupiter* fluorescence is coded with the rainbow spectrum shown (A.U.). Scaled axes are provided in µm. (**C**) Heat map representing the average normalized, binned *mCherry::Jupiter* fluorescence along dendrites at increasing distances from the soma for each genotype. TORC1 inhibitions are marked in purple and TORC1 activations in green. Genotypes found to be significantly different along the dendritic arbor are marked with an asterisk. In all panels * = p < 0.05, see **Supplementary Table S2** for detailed statistics.

We examined the levels of Futsch – a microtubule-associated protein (MAP), acetylated α-tubulin, and βtubulin IIA in the soma of CIV neurons. Both Futsch and acetylated α-tubulin serve as markers of stable MTs (Hummel et al., 2000; Pawson et al., 2008; Weiner et al., 2016; Eshun-Wilson et al., 2019), and β-tubulin IIA is a MT subunit known to be specifically folded by CCT (Llorca, 2000).

In CIV neurons, basal levels of acetylated α-tubulin are significantly reduced in *CCT3-IR*, *CCT5-IR*, *Raptor-IR*, *S6k-IR*, and *Akt-IR*, but are not significantly changed in *Cullin1-IR, S6k OE,* or *Akt OE* (**Fig 3A**). Futsch is significantly decreased in TORC1 inhibition conditions: LOF of *CCT3*, *CCT5*, *S6k*, *Raptor*, or *Akt* leads to significant reductions of Futsch fluorescence (**Fig 3A**). In contrast, *Akt* and *S6k* overexpression leads to significant increases in Futsch signal (**Fig 3A**); however, *Cullin1-IR* does not show a significant change from control.

*CCT3* and *CCT5* LOF lead to significant decreases in β-tubulin IIA, as CCT LOF does for measures of MT stability (**Fig 3A, S2E**). Although *Akt* and *Raptor* LOF also significantly reduce β-tubulin IIA fluorescence, surprisingly, *S6k* LOF significantly increases overall levels of β-tubulin IIA compared to WT neurons. Interestingly, *Cullin1* also significantly decreases β-tubulin IIA fluorescence (**Fig S2F**).

We further confirm that TORC1 LOF reduces stable MT levels throughout the dendritic arbor through the use of a fluorescent line marking the MT-associated protein Jupiter (*UAS-Jupiter::mCherry*) (Das et al., 2017). CCT LOF (1-, 3-, and 5-IR) results in the steepest decline in *Jupiter::mCherry* signal (**Fig 3B-C**). Similar to the effects on acetylated α-tubulin and Futsch, loss of *Raptor*, *S6k*, and *Akt* also significantly decrease *Jupiter::mCherry* signal; however, *Akt* OE and *S6k* OE do not significantly alter *Jupiter::mCherry* signal. Interestingly, *Cullin1* knockdown significantly decreases *Jupiter::mCherry* signal despite also resulting in hyper-arborization (**Fig 3B-C**). In general, we find that TORC1 inhibition significantly decreases MT signal along the dendritic arbor, and that TORC1 hyperactivation results in varied MT signal along the arbor depending on the measured marker of MT stability.

### Mutant Huntingtin expression leads to repeat length-dependent reduction in branch complexity and underlying microtubule losses

Though CCT and TORC1 clearly regulate dendritic arborization during development in homeostatic conditions, there is also great interest in examining the putative relationships of these complexes to proteinopathic disease. Several studies have connected CCT to the regulation of mutant Huntingtin (mHTT) protein, and there is some evidence that loss of wild-type HTT disrupts neuron formation (McKinstry et al., 2014; Barnat et al., 2017). *Drosophila melanogaster* has been used to model many HD-related phenomena, such as motor deficits, circadian rhythm changes, metabolic precursors of disease, mHTT aggregate spreading in the brain, and more (Krench and Littleton, 2013; Bertrand et al., 2020; Vernizzi et al., 2020; Khyati et al., 2021; Subhan and Siddique, 2021). Using UAS-mediated constructs of mutant Huntingtin (see **Supplementary Table S1**), we expressed human mHTT in CIV neurons and quantified gross morphology of the resultant dendritic arbors. Shorter repeats of mHT and mHT do not significantly alter arbor complexity, but expression of mHT and mHT0 significantly reduce TDL from WT (**Fig S3A-B**). We also examined HTT distribution in CIV neurons and identified somatic and dendritic expression (**Fig 4A**). Consistent with WT HTT expression, induced expression of mHTT120HA and mHTT96Cer show clear expression in soma and dendrites (**Fig 4B, S3E**). Though mHTT misexpression lines have significantly higher overall HTT expression than WT (**Fig S3G**), neurons expressing *UAS-mHTTQ25-Cerulean* do not display apparent puncta (**Fig 4B**). In contrast, misexpression of *UAS-mHTTQ96-Cerulean* or *UAS-mHTTQ120-HA* results in aggregate inclusion bodies (IBs) of mHTT forming in the dendritic arbor (**Fig 4B, S3E**).

**Figure 4:**
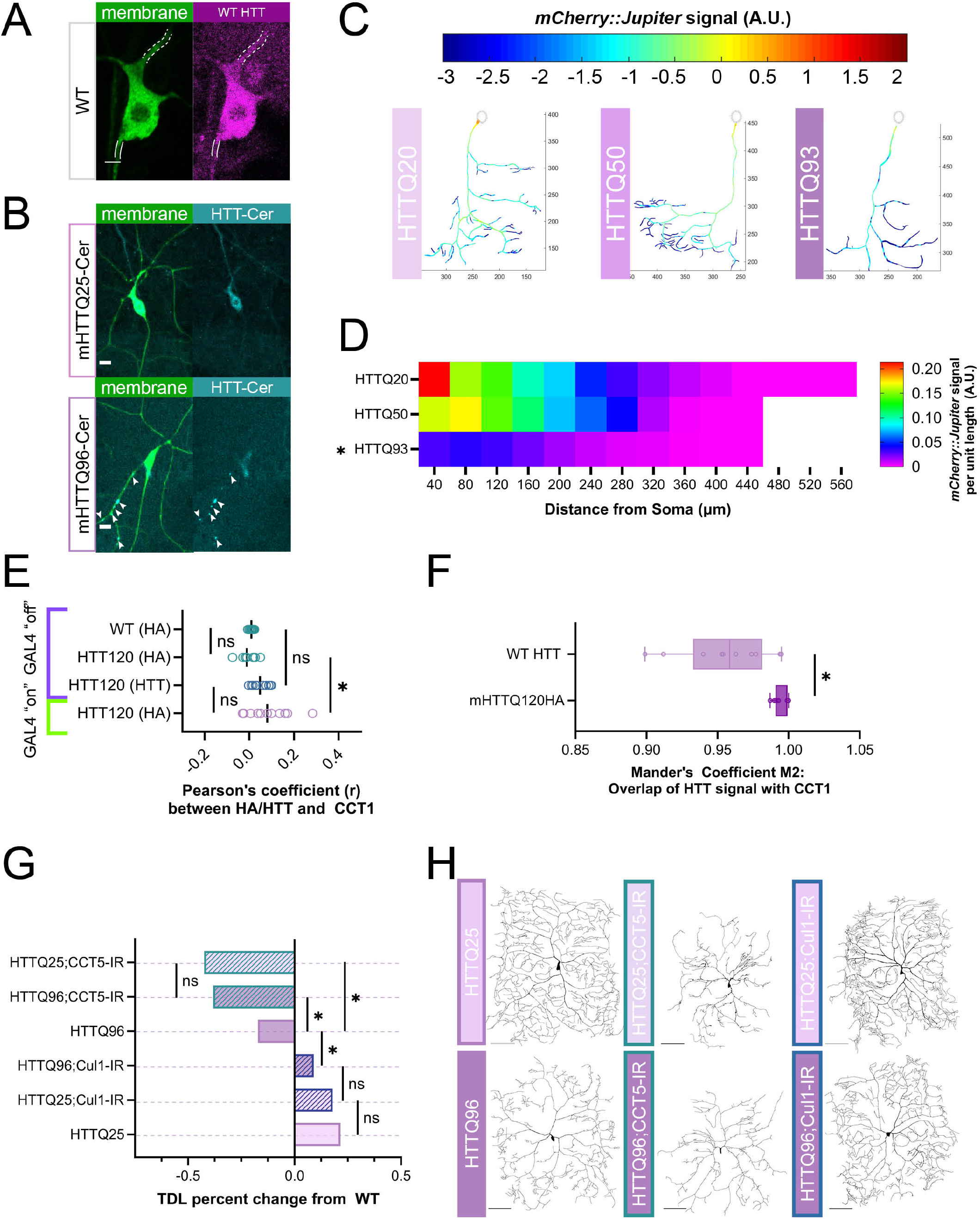
Expression of mHTT leads to dendritic hypotrophy parallel to TORC1 pathway. (**A**) Representative image of WT HTT staining in CIV neuron – dendrite marked by dashed white lines, axon by solid lines. Scale bar = 3 µm. (**B**) Representative images of mHTT25-Cerulean and mHTT96-Cerulean shown with aggregate inclusion bodies marked by white arrows for mHTT96-Cerulean. Scale bar = 10 µm. (**C**) Representative reconstructions of branches from WT and TORC1 genetic manipulations – normalized *mCherry::Jupiter* fluorescence is coded with the rainbow spectrum shown (A.U.) (**D**) Heat map representing the average normalized, binned *mCherry::Jupiter* fluorescence along dendrites at increasing distances from the soma for overexpressions of mHTT 20, 50, and 93 repeats. Genotypes found to be significantly different along the dendritic arbor are marked with an asterisk. (**E**) Pearson’s correlation Coefficient (*r*) is near-zero for CCT1 and HTT co-expression in soma cytosol, however, *r* for CCT1-mHTT120HA is significantly higher than CCT1-WT HTT and HA-stained Gal4 “off” controls. (**F**) Thresholded Mander’s Coefficient M2 signifying overlap of two signals is significantly higher in CCT1-mHTT120HA conditions than CCT1-WT HTT conditions. (**G**) TDL of *Cullin1-IR* and *CCT5-IR* in both *mHTTQ25* and *mHTTQ96* backgrounds displayed as percent change from WT control. *CCT5-IR* decreases both mHT and mHT neurons to far lower than WT, while *Cullin1-IR* rescues mHTT96 hypotrophy to WT levels. (**H**) Representative images of CIV dendritic morphology in combined *HTT* and *CCT5-IR* or *Cullin1-IR* combinations. Scale bars = 100 µm. In all panels * = p < 0.05, see **Supplementary Table S2** for detailed statistics.

WT HTT is thought to be involved in cellular trafficking, and there is evidence that mHTT expression can destabilize MTs (Trushina et al., 2003; Subhan and Siddique, 2021). We predicted that there would be underlying cytoskeletal deficits in these neurons similar to the phenotypes we observe in CCT and TORC1 LOF neurons. Expression of *mHTTQ96* significantly decreases Futsch fluorescence levels in the soma as measured via IHC (**Fig S3C**). Live imaging of *Jupiter::mCherry,* reveals that expression of *mHTTQ50* does not significantly reduce stable MT signal, but expression of *mHTTQ93* significantly reduces stable MT signal across the arbor (**Fig 4C-D**) as compared to the non-phenotypic mHT. Overall, expression of high repeats of mHTT results in dendritic hypotrophy and underlying losses of stable MTs in CIV neurons.

### mHTT induction of dendritic hypotrophy may involve CCT and TORC1

A large number of previous studies implicate CCT in direct regulation of mHTT (Tam et al., 2006; Shahmoradian et al., 2013; Sontag et al., 2013; Noormohammadi et al., 2016; Shen et al., 2016; Zhao et al., 2016); therefore, we first sought to answer whether CCT regulates wild-type Huntingtin. We found that *CCT5* knockdown led to a significant decrease in soma levels of wild-type *Drosophila* Huntingtin, as measured through IHC (**Fig S3D**).

CCT has been shown to reduce aggregation of mutant Huntingtin (mHTT) through physical interaction *in vitro* (Tam et al., 2006; Shahmoradian et al., 2013), thus we predicted that subunits of CCT would co-localize with HTT and mHTT in the cytosol. To test this prediction, we examined the correlation of expression between CCT1 and HTT, as well as between CCT1 and mHT0-HA (**Fig 4E-F, S3E**). The use of the temperature-sensitive Gal4 inhibitor, *ts-Gal80*, allowed us to measure HA and HTT levels in WT neurons and those with mHT0-HA expressed (Gal4 “on”) as well as those with mHT0-HA suppressed (Gal4 “off”) (See Methods for details) (Fig S3E).

Analyses of CIV neuron soma from these genetic backgrounds reveal near-zero Pearson’s correlation Coefficients (PC) between CCT1 and WT HTT as well as between CCT1 and mHT0-HA (**Fig 4E**). PC is a measurement of the linear relationship between two fluorescent labels in an area of interest that ranges from -1 (perfect exclusion) to 1 (perfect colocalization) (Cordelières and Bolte, 2014). We find a small, but significant increase in the correlation for mHTT120HA from its positive genetic control but no difference between mHTT120HA and WT HTT (**Fig 4E**). Since PC is sensitive to noise and non-linear relationships between labels, we also analyzed the images using Mander’s correlation Coefficient, with strict predetermined thresholds (see Methods for details). The M2 statistic, which describes the fraction of the CCT1 signal overlapping with HTT, was high in both WT and mHTT conditions (**Fig 4F**). There was a small, but significant increase in the M2 statistic between WT and mHTT conditions in both soma and dendrites (**Fig 4F, S3F**). In sum, CCT1 and both WT HTT and mHTT120-HA are clearly present in the cytoplasm of CIV soma and dendrites, and though they display high overlap, we find no linear relationship between the two signals at the current resolution.

As CCT appears to be necessary for WT HTT expression, and the two proteins show high co-expression in the cytoplasm, we predicted that they might operate within the same pathway to regulate dendritic arborization. After knocking down *CCT5* in neurons expressing either subclinical *mHTTQ25-Cerulean* or *mHTTQ96-Cerulean*, we find the combination does not show potentiation of the two phenotypes. *CCT5-IR* appears to strongly drive the hypotrophy phenotype in both mHT and mHT backgrounds (**Fig 4G-H**).

Previous *in vitro* evidence suggests that CCT may work to clear mHTT aggregates (Kitamura et al., 2006; Tam et al., 2006; Shahmoradian et al., 2013; Sontag et al., 2013; Sergeeva et al., 2014; Shen et al., 2016), thus we predicted that CCT LOF may lead to higher aggregate load in dendrites *in vivo*. However, CCT LOF in the *UAS-mHTTQ96-Cerulean* background does not lead to a significant change in IB number or size (**Fig S3H-I**). Additionally, *CCT5-IR* in a *mHTT96* background does not potentiate the loss of Futsch signal, which is already significantly reduced in mHT neurons (**Fig S3C**). Though CCT is required for WT HTT expression and shows high co-expression with both WT HTT and mHTT in the cytosol, CCT LOF does not appear to exacerbate IB appearance at the current resolution.

TORC1 activity has also been explored as an avenue for mHTT clearance, mainly through inhibition via the application of rapamycin (Sarkar et al., 2008; Pryor et al., 2014; Lee et al., 2015). Given HTT’s many connections to both CCT and TORC1, we predicted that expression of mHTT may influence dendritic arborization through disruption of the insulin pathway and cytosolic interactors CCT and Cullin1. Interestingly, knockdown of *Cullin1* in the *mHTTQ96* background increases TDL significantly from *mHTT96* alone, returning it to WT levels (**Fig 4G-H**). Despite rescue of the dendritic hypotrophy, when *Cullin1* is knocked down in a *UAS-mHTTQ96-Cerulean* background, neither the median number, nor aggregate size, of mHTT IBs changes (**Fig S3H-I**). Overall, though we find that CCT5 is required for WT HTT levels, and that CCT1 and HTT, as well as mHTT, are both co-expressed in the cytoplasm of CIV soma and dendrites, we did not find that CCT LOF and mHTT expression resulted in an additive phenotype. Interestingly, we did find that *Cullin1* LOF was sufficient to rescue mHT-mediated defects in TDL, though it did not significantly affect the appearance of mHTT dendritic IBs.

## DISCUSSION

### A TORC1 cytosolic network regulates dendritic development and the underlying MT cytoskeleton

The TORC1 pathway has many cytosolic interactors; we have illuminated the roles of two, CCT and Cullin1, in dendritic development. Previous work established that CCT subunits are required for CIV dendritic arbor formation (Das et al., 2017; Wang et al., 2020), and we have confirmed that CCT LOF results in dendritic hypotrophy with underlying stable MT deficits. Though CCT directly folds actin and tubulin monomers, we predicted that its contribution to dendritic arborization may also extend to secondary regulators of the dendritic arbor, such as TORC1. TORC1 has been found to regulate dendritic arbors in mammalian dopaminergic neurons (Diaz-Ruiz et al., 2009; Kosillo et al., 2019, 2022), and was recently found to be regulated by CCT in both *Drosophila* and human cell cultures (Vinayagram et al., 2016; Cuéllar et al., 2019; Kim and Choi, 2019). We confirmed, *in vivo,* that CCT is required for WT levels of Raptor and P-S6k in *Drosophila* CIV neurons. *CCT5* knockdown significantly reduces Raptor expression levels and cannot be rescued by Raptor overexpression, indicating that CCT is required for WT levels of Raptor (**Fig 2B**). Additionally, S6k overexpression could not rescue P-S6k levels in *CCT5* knockdown neurons, indicating that CCT is required for WT levels of P-S6k (**Fig 2C**).

Furthermore, we found TORC1 LOF results in dendritic hypotrophy while TORC1 hyperactivation results in dendritic hypertrophy (**Fig 1B-D**). The hypotrophy resulting from both LOF of TORC1 and CCT is mirrored by underlying losses of stable MT markers in the soma such as Futsch and acetylated α-tubulin as well as *Jupiter::mCherry* signal throughout the dendritic arbor (**Fig 3A-C**). A notable difference is that CCT, Raptor, and Akt LOF significantly reduce β-tubulin IIA signal in the soma, while S6k LOF significantly increases β-tubulin IIA signal (**Fig S2E**). The β-tubulin IIA antibody used in our experiments is not specific to either free or incorporated β-tubulin IIA, so the production of free β-tubulin IIA could create increased fluorescence even in a cell with reduced stable MTs.

Our gross morphological findings of TORC1 LOF coincide tightly with those of a recent study demonstrating that changes in nutrition result in CIV dendritic hyper-arborization and subsequent changes in cell sensitivity and larval behavior (Kanaoka et al., 2023). Akt, TOR, and S6k were all found to be required for hyper-arborization induced by a low-yeast diet. Levels of phosphorylated Akt were increased in low-yeast diet conditions, and overexpression of Akt increased dendritic complexity. *Akt* LOF and OE have been established to decrease and increase CIV dendritic coverage, respectively (Parrish et al., 2009). In our work, LOF of Akt and S6k produce dendritic hypotrophy (**Fig1B-D**), and while we have found that there are underlying MT deficits in TORC1 LOF conditions (**Fig 3A-C**), it remains to be seen if the cytoskeletal phenotypes are inducible through diet changes.

Manipulations inducing TORC1 hyperactivity or disinhibition of TORC1 all result in hypertrophy but have variable effects on MT markers. In our experiments, inhibition of TORC1 pathway genes reduces levels of acetylated α-tubulin while TORC1 hyperactivity leads to increases in acetylated α-tubulin by 15-22% (**Fig 3A**). S6k has previously been found necessary for stress-evoked acetylation of α-tubulin in mouse embryonic fibroblasts (Hać et al., 2021), but the mechanism connecting S6k to tubulin acetylation is still unclear. TORC1 hyperactivation through *S6k* OE and *Akt* OE also increases Futsch levels; however, *Cullin1-IR*-mediated disinhibition of TORC1 does not show the same MT phenotypes despite similar dendritic hypertrophy (**Fig 3A-C**). Although *Cullin1-IR* results in a 22% increase in acetylated α-tubulin, it results in a mild decrease in Futsch and a significant decrease in Jupiter signal. It may be that loss of *Cullin1* reduces the expression or attachment of MAPs to MTs without affecting underlying MT stability. Together, our data suggest that CCT and Cullin1 partially influence dendritic arborization and development of the neuronal cytoskeleton through regulation of TORC1.

### TORC1 cytosolic network and mHTT interact in the regulation of dendritic arbors

Although important to understand the role of TORC1 in homeostatic dendritic development, TORC1 has also been extensively investigated with respect to proteinopathies, including Huntington’s Disease (for recent review see (Querfurth and Lee, 2021)) as has CCT (Tam et al., 2006; Shahmoradian et al., 2013; Sergeeva et al., 2014; Noormohammadi et al., 2016; Shen et al., 2016; Zhao et al., 2016; Chen et al., 2018). Similar to TORC1 and CCT LOF phenotypes, expression of high repeat human mHTT in CIV neurons reduces dendritic complexity and underlying stable MT signal (**Fig S3 A-B, 4C-D, G-H**). We observed both WT and mutant HTT in multiple neuronal compartments; furthermore, expression of mHT-Cerulean and mHT-HA produced large aggregate IBs in dendritic arbors, similar to those previously reported in (Krench and Littleton, 2013) (**Fig 4A-B, S3E**).

Although CCT appears to be required for WT levels of HTT, we found mixed colocalization results for CCT1 and HTT (**Fig 4E-F, S3F**). Based on the current level of resolution, we find no linear relationship between CCT1 and HTT signal in the cytoplasm despite a high degree of co-expression in both soma and dendritic compartments (**Fig 4E-F, S3F**). The correlation between CCT1 and mHTT120HA was significantly higher than the correlation between CCT1 and HTT; however, as all conditions are still far closer to zero than one, we do not assert that there is any meaningful correlation between CCT1 and WT HTT or mHTT120-HA signal within the cytoplasm. Given the differences between the two statistics we examined (PC and Mander’s), it is possible that HTT and CCT1 have a non-linear co-expression relationship in the cytoplasm (*e.g.* CCT1 signal could increase near mHTT aggregates but not at the rate of mHTT signal increase).

We also carried out genetic interaction studies between CCT and mHTT expression. When combined, *CCT5* LOF and *mHTTQ96* expression do not display an additive dendritic phenotype: arbor complexity in the mHTT96 background is reduced to the same level as that in neurons with both mHTT25 and *CCT5-IR* (**Fig 4G-H**). Furthermore, *CCT5-IR* did not induce a change in size or number of mHTT dendritic puncta in the *mHTTQ96-Cerulean* background (**Fig S3H-I**). This is in contrast with previously published iPSC data, which found LOF of individual CCT subunits triggered aggregation of mHTT (Noormohammadi et al., 2016). CIV neurons form IBs upon expression of high repeat mHTT, whereas iPSCs expressing mHTT require heat or proteostatic stress to induce IB formation, which may explain the discrepancy. There may also be changes in the IBs that are not evident at our current experimental parameters, such as organization of mHTT within the IB, or changes to temporal dynamics of IB formation.

Cullin1, the cytosolic inhibitor of TORC1, does show genetic interaction with mHTT expression: when we knocked down *Cullin1* in *mHTTQ96* neurons, there was a significant increase in dendritic complexity, restoring TDL to WT levels (**Fig 4G-H**). Unexpectedly, in this same genetic background, we did not observe any change to dendritic IB size or number (**Fig S3H-I**). We had predicted *Cullin-IR* would lead to an increase in IB number or size for two reasons. First, Cullin1, as part of an E3 ubiquitin ligase, helps to ubiquitinate proteins for degradation (Gerez et al., 2019), and *Cullin1* LOF has been linked to increased protein aggregate load (Bhutani et al., 2012; Chen et al., 2019). Second, *Cullin1-IR* leads to TORC1 disinhibition, which reduces autophagic activity (Switon et al., 2017). Previous studies have shown that rapamycin application – inhibition of TORC1 – reduces mHTT aggregation in *Drosophila* ommatidia and mammalian cells (Ravikumar et al., 2004; Berger et al., 2006), and other studies have shown that inhibition of mTORC1 ameliorates mHTT pathology through increased autophagic activity (Berger et al., 2006; Roscic et al., 2011; Vernizzi et al., 2020; Querfurth and Lee, 2021). However, we find in CIV neurons, *Cullin1* knockdown increases dendritic complexity while the dendritic IB load remains unchanged. There is evidence that the SCF complex is down-regulated in Parkinson’s Disease, Huntington’s Disease, and Spinal-Cerebellar Ataxia Type 3, and that further *Cullin1* LOF exacerbates aggregate phenotypes (Bhutani et al., 2012; Mandel et al., 2012a, 2012b; Chen et al., 2019). Therefore, it is possible that the role of Cullin1 in suppression of complex dendritic development and its role in promoting degradation of protein aggregates are carried out through distinct cellular pathways. There are undoubtedly several mechanisms by which complexes like TORC1, CCT, and SCF could interact with protein aggregates, providing fertile ground for future studies.

In summary, our study shows that CCT regulates TORC1 *in vivo* to promote dendritic arborization in homeostatic development. We further demonstrate that Cullin1 inhibits TORC1 *in vivo* to suppress dendritic arborization. At the cytoskeletal level, TORC1 hypo-activation leads to underlying stable MT deficits, while hyper-activation and disinhibition through *Cullin1* knockdown have distinct MT phenotypes. In proteinopathic disease conditions, high repeats of mHTT lead to dendritic hypotrophy, and *Cullin1* LOF can rescue mHTT-induced hypotrophy, though neither *Cullin1* LOF nor *CCT* LOF significantly alters mHTT aggregate IB expression. Our data, together with previous literature, demonstrate conserved roles of TORC1, CCT, and Cullin1 in dendritic regulation in healthy and diseased neurons.

## ACKNOWLEDGEMENTS

This work is supported by NIH R01 NS086082 to DNC. ENL was supported by an NIH F31 NS130970-01, a GSU Brains and Behavior (B&B) Fellowship, a GSU 2CI Neurogenomics Fellowship, and a Kenneth W. and Georganne F. Honeycutt Fellowship. FHC was supported by a GSU 2CI Neurogenomics Fellowship. EAT-W and TT were supported by NIH Maximizing Access to Research Careers (MARC) fellowships (T34 GM131939-01). BT was supported by a GSU B&B summer fellowship. AAP was supported by a GSU 2CI Neurogenomics Fellowship, a Kenneth W. and Georganne F. Honeycutt Fellowship, and a B&B Fellowship. We thank Dr. Kwang-Wook Choi at KAIST, South Korea for the *UAS-dCCT4* transgenic line. The authors gratefully acknowledge the Imaging Core Facility at Georgia State University for their support and assistance in this work.

ENL *and* DNC designed the research. ENL, FHC, SB, and EAT-W performed the research. ENL, FHC, SB, EAT-W, BT, TT, and JM analyzed the data. AAP contributed analytic tools. ENL wrote the manuscript, which was revised by FHC, SB, and DNC. ENL and DNC were responsible for funding acquisition and DNC for project supervision.

## Supplementary Figures

**Figure S1:**
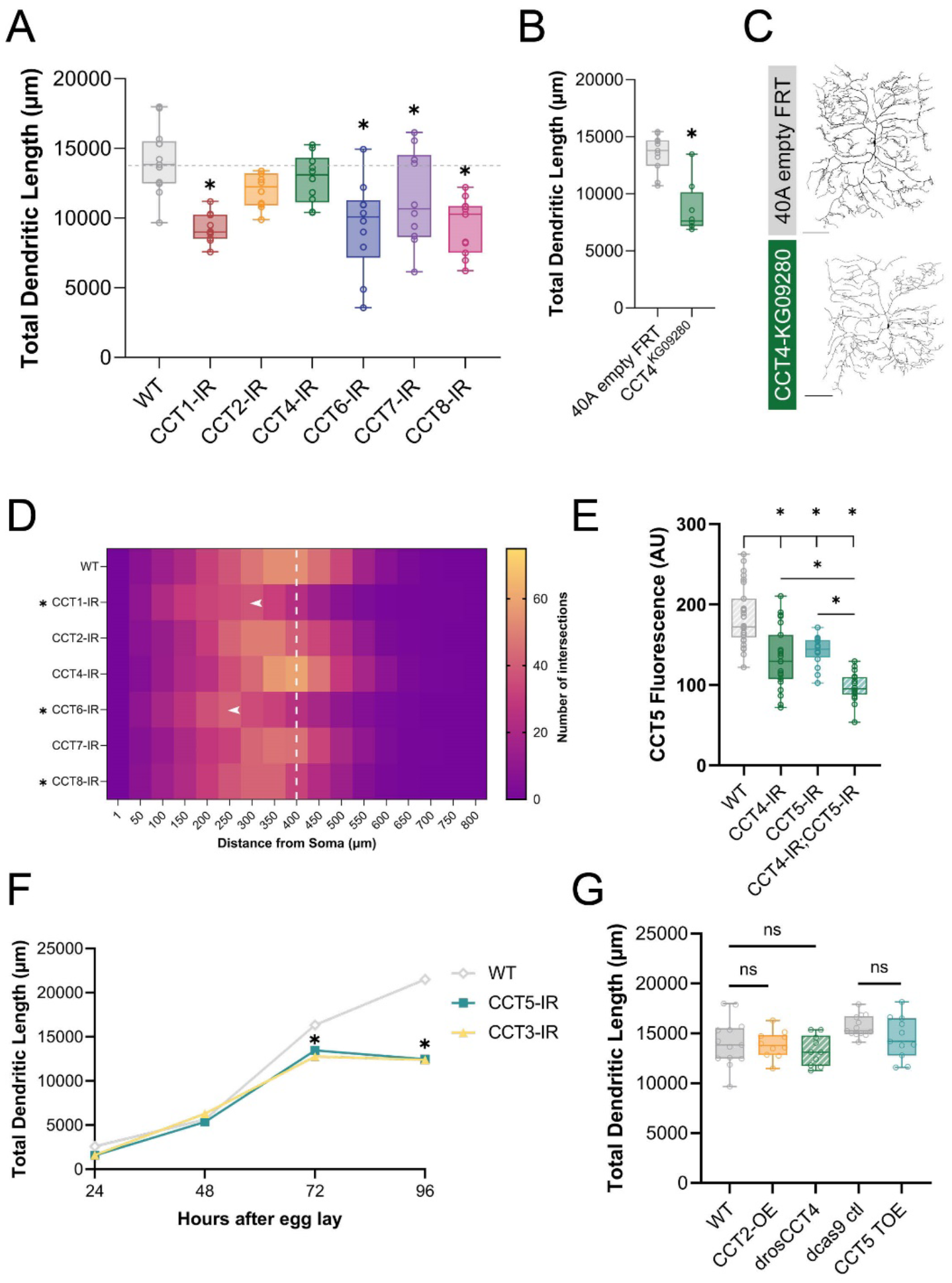
CCT subunit LOF results in significant hypotrophy and underlying loss of stable MTs. (**A**) Loss of individual CCT subunits results in significant decreases in TDL from WT controls. (**B**) Homozygous CCT4 MARCM mutant clones show significantly decreased TDL from control. (**C**) Representative images of CCT4 homozygous MARCM mutant CIV clones vs. control CIV MARCM clones (40A empty FRT). (**D**) Number of Sholl intersections mapped by color at increasing radial distances from soma (µm). White dashed line references the maximum Sholl intersections in WT neurons. Significant changes in Sholl maximum intersections are indicated by an asterisk. Arrows indicate genotypes where the radius of maximum intersections has shifted significantly from WT. (**E**) RNAi of *CCT4* or *CCT5* lead to a significant reduction in CCT5 fluorescence relative to WT as obtained through IHC. Combined knockdown of both *CCT4* and *CCT5* significantly reduces CCT5 expression from either knockdown alone. (**F**) TDL of neurons at 24, 48, 72, and 96 hours after egg lay (AEL) reveal significant decreases from WT in both *CCT5-IR* and *CCT3-IR* starting at 72 hours AEL. (**G**) Overexpression of individual CCT subunits (CCT2, CCT4, or CCT5) does not significantly alter TDL from their relevant control. In all panels * = p < 0.05, see **Supplementary Table S2** for detailed statistics.

**Figure S2:**
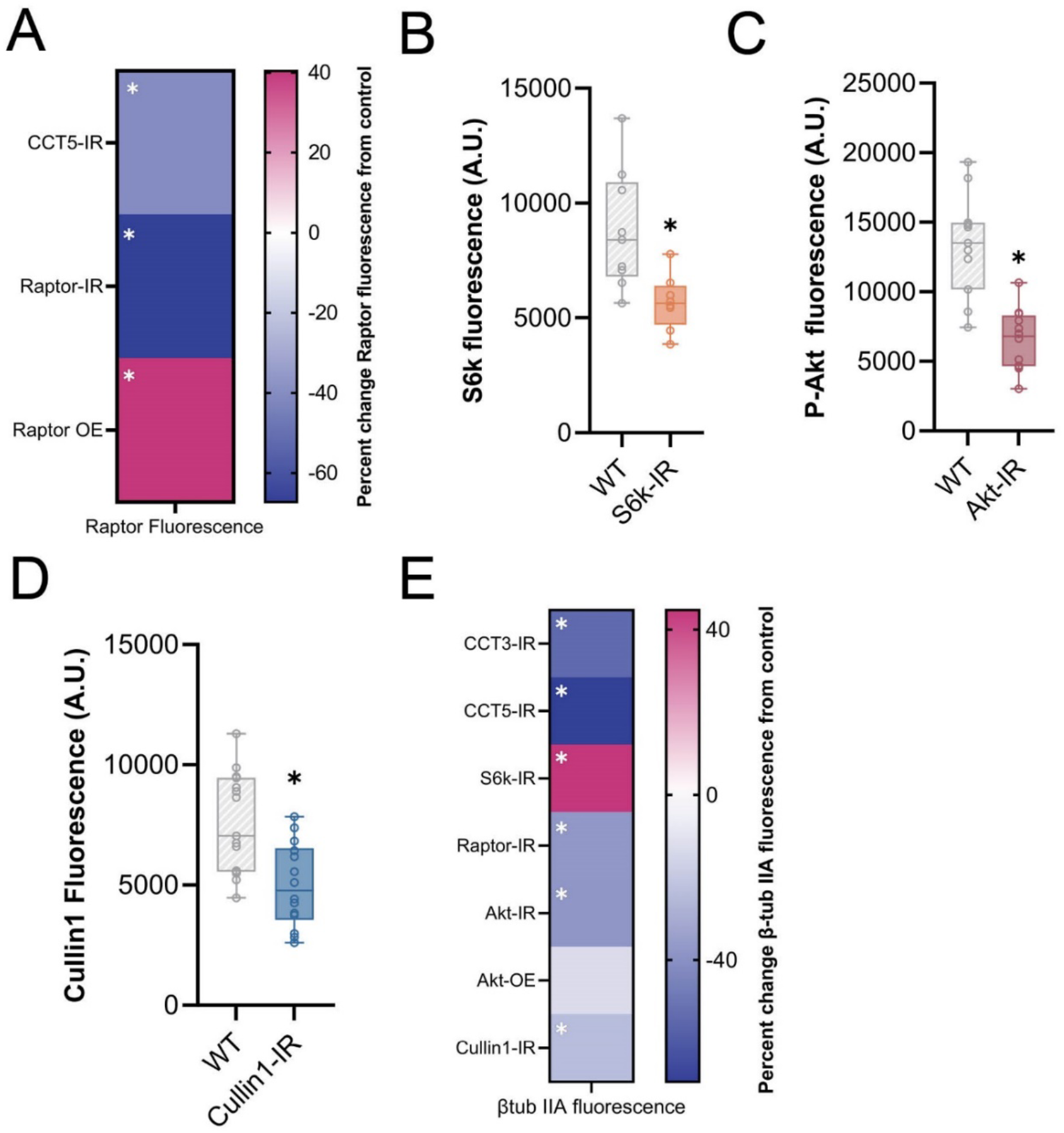
Evidence for RNAi efficacy and CCT and Cullin1 regulate the TORC1 pathway *in vivo*. (**A**) Heat map showing percent change in Raptor fluorescence of CCT5-IR or Raptor-IR knockdowns, as well as Raptor OE as compared to controls. (**B**) S6k fluorescence is significantly reduced in *S6k-IR* conditions as compared to WT. (**C**) P-Akt fluorescence is significantly reduced in *Akt-IR* conditions. (**D**) Cullin1 fluorescence is significantly reduced in *Cullin1-IR* conditions as compared to WT. (**E**) Heat map showing percent change in β -tubulin IIA for each genetic manipulation. Each experimental condition was compared to WT control and appropriate statistical comparisons were performed. In all panels * = p < 0.05, see **Supplementary Table S2** for detailed statistics.

**Figure S3:**
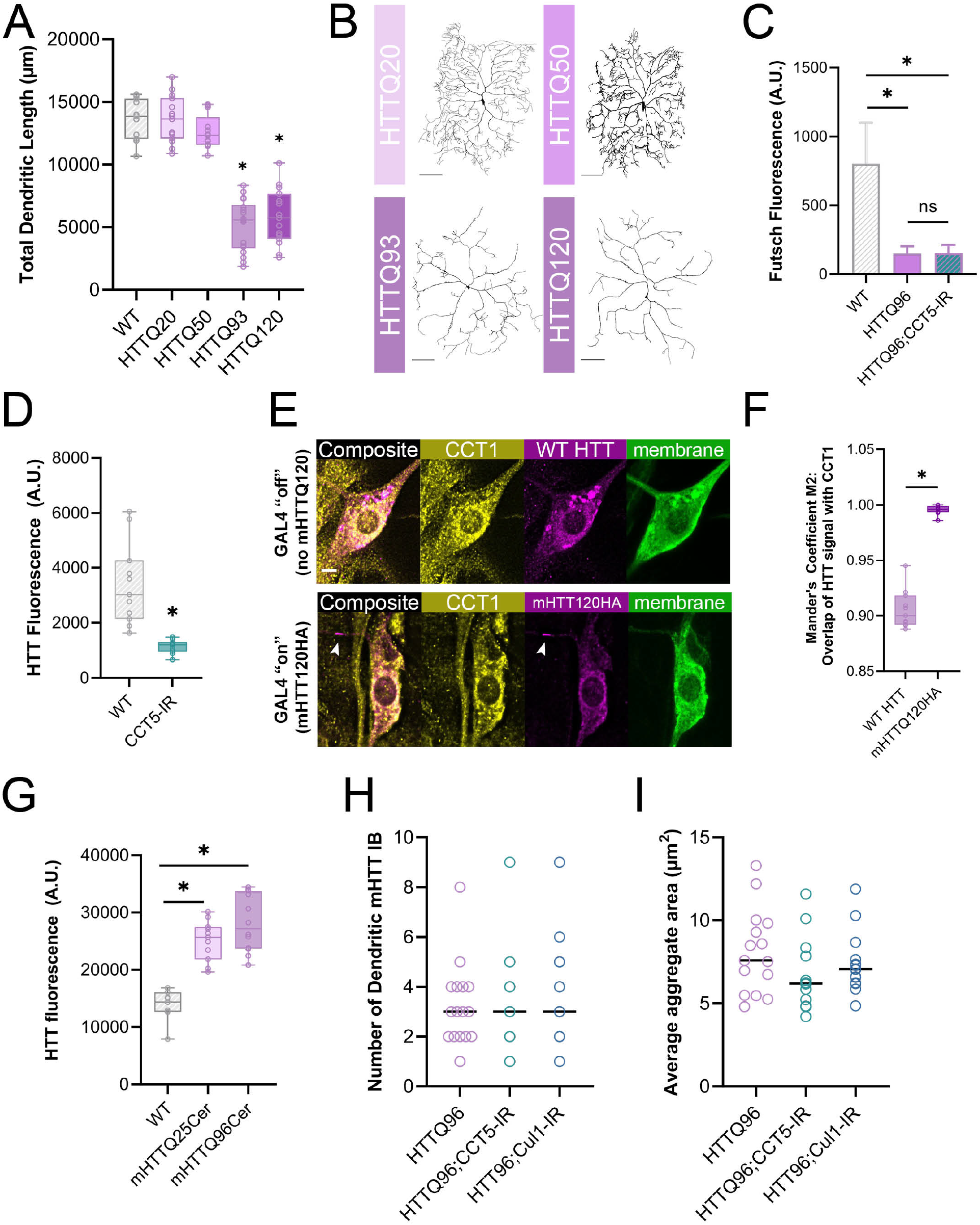
mHTT aggregates are not affected by genetic combinations despite high co-expression of CCT1 and HTT. (**A**) TDL is significantly reduced from control in neurons expressing mHTT93 or mHTT120 CAG repeats. (**B**) Representative images of CIV neurons expressing mHTT polyQ repeat transgenes reveal repeat-length dependent dendritic hypotrophy. Scale bars = 100 µm. (**C**) Fluorescent levels of Futsch are significantly reduced in mHT conditions and are not significantly changed by additional *CCT5-IR* expression. (**D**) WT HTT fluorescence is significantly reduced in *CCT5* LOF conditions. (**E**) Representative images of WT HTT distribution in mHTT120HA suppressed (Gal4 “off”) and mHTT120HA distribution in Gal4 “on” conditions. Aggregate IB indicated by arrow in dendrite. Scale bar = 3 µm. (**F**) Mander’s M2 coefficient is significantly increased for co-expression of CCT1 and mHTT120HA as compared to CCT1 and WT HTT in dendrites. (**G**) Expression of mHTT25-Cerulean or mHTT96-Cerulean both result in significant increases in HTT fluorescence from WT. (**H**) Number of mHTT aggregate IBs does not change due to *CCT5* or *Cullin1* LOF. (**I**) mHTT aggregates in mHTCerulean conditions do not change in average area due to *CCT5* or *Cullin1* LOF. In all panels * = p < 0.05, see **Supplementary Table S2** for detailed statistics.

**Supplementary Genetics Table S1.**
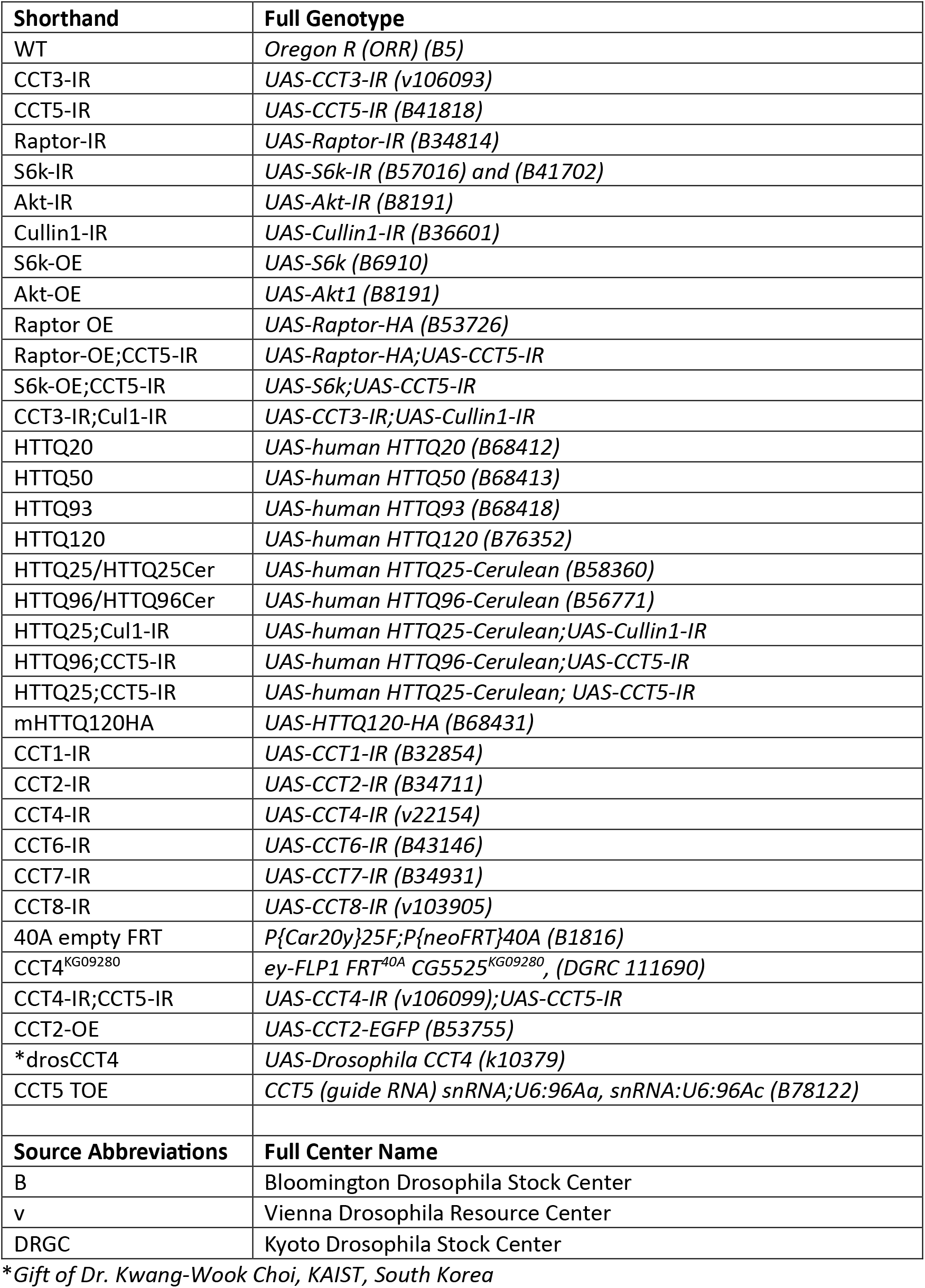
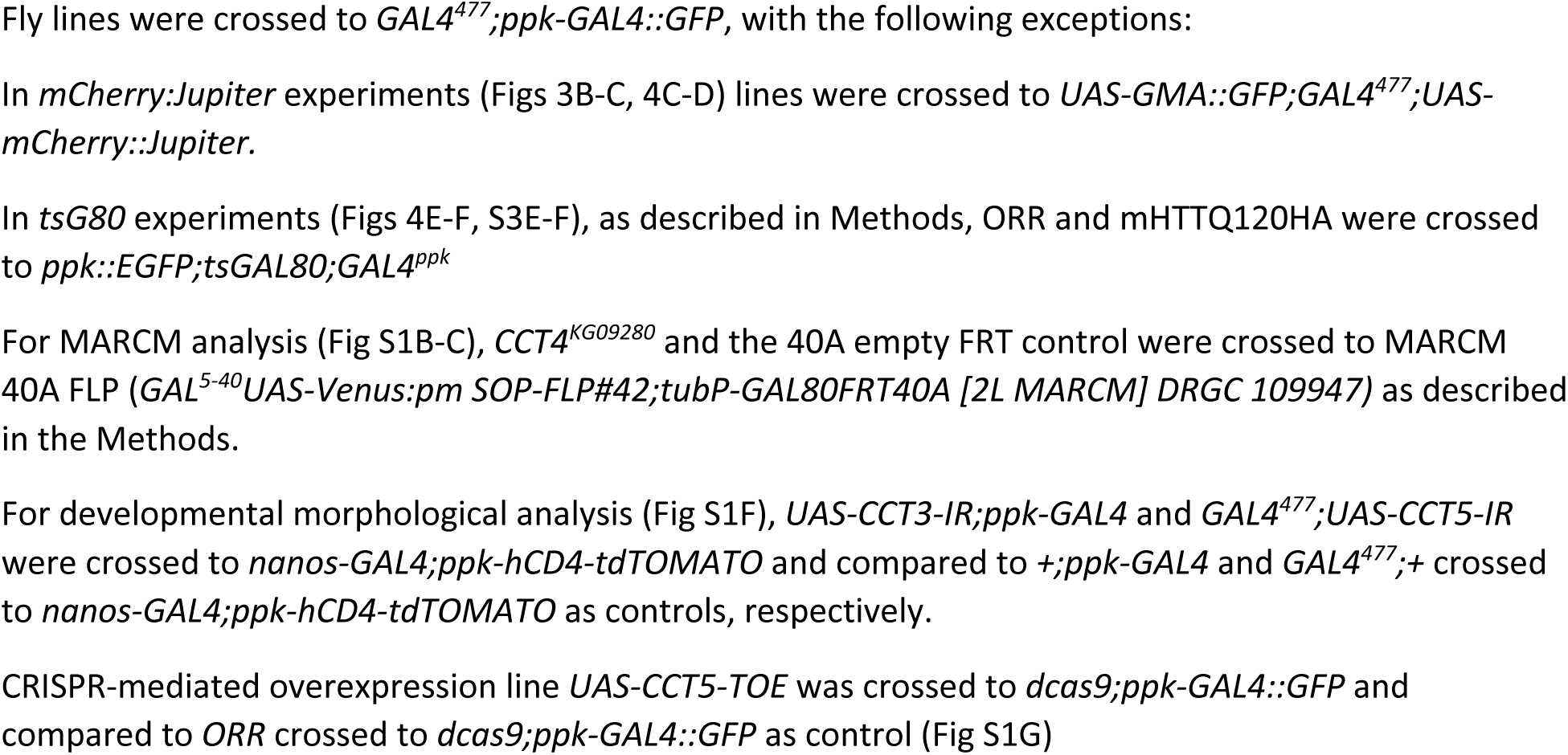

**Supplementary Statistics Table S2.**

| Comparison | Passed Shapiro-Wilk Normality Test | Statistical test used | Sig | p-value | Number of neurons (N) |
| --- | --- | --- | --- | --- | --- |
| **Fig 1C (TDL)** |  | One-way ANOVA & Dunnett's |  |  |  |
| WT vs. CCT3-IR | Yes |  | **** | <0.0001 | 13, 11 |
| WT vs. CCT5-IR | Yes |  | **** | <0.0001 | 13, 12 |
| WT vs. Raptor-IR | Yes |  | *** | 0.0002 | 13, 10 |
| WT vs. S6k-IR | Yes |  | **** | <0.0001 | 13, 13 |
| WT vs. Akt-IR | Yes |  | **** | <0.0001 | 13, 10 |
| WT vs. Cul1-IR | Yes |  | *** | 0.0003 | 13, 13 |
| WT vs. S6k-OE | Yes |  | **** | <0.0001 | 13, 10 |
| WT vs. Akt-OE | Yes |  | **** | <0.0001 | 13, 11 |
| **Fig 1D (Sholl Maximum)** |  | One-way ANOVA & Dunnett's |  |  |  |
| WT vs. CCT3-IR | Yes |  | * | 0.0219 | 9, 11 |
| WT vs. CCT5-IR | Yes |  | ns | >0.9999 | 9, 14 |
| WT vs. S6k-IR | Yes |  | ns | 0.2517 | 9, 14 |
| WT vs. Akt-IR | Yes |  | **** | <0.0001 | 9, 10 |
| WT vs. Akt-OE | Yes |  | ** | 0.0012 | 9, 11 |
| WT vs. S6k-OE | Yes |  | **** | <0.0001 | 9, 10 |
| WT vs. Cul1-IR | Yes |  | ** | 0.0027 | 9, 12 |
| **Fig 1D (Sholl Radius)** |  | One-way ANOVA & Dunnett's |  |  |  |
| WT vs. CCT3-IR | Yes |  | ns | 0.0522 | 9, 11 |
| WT vs. CCT5-IR | Yes |  | **** | <0.0001 | 9, 14 |
| WT vs. S6k-IR | Yes |  | ns | 0.9261 | 9, 14 |
| WT vs. Akt-IR | Yes |  | ns | 0.7088 | 9, 10 |
| WT vs. Akt-OE | Yes |  | * | 0.0126 | 9, 11 |
| WT vs. S6k-OE | Yes |  | **** | <0.0001 | 9, 10 |
| WT vs. Cul1-IR | Yes |  | ns | 0.9965 | 9, 12 |
| **Fig 2A (P-S6k IHC)** |  |  |  |  |  |
| WT vs. S6k-IR | No | Mann-Whitney test | **** | <0.0001 | 14, 12 |
| WT vs. CCT5-IR | Yes | One-way ANOVA & Tukey's | *** | 0.0008 | 9, 9 |
| WT vs. Raptor-IR | Yes | One-way ANOVA & Dunnett's | ** | 0.0099 | 14, 9 |
| WT vs. Raptor-OE | Yes | One-way ANOVA & Dunnett's | ns | 0.9612 | 14, 16 |
| WT vs. Cullin1-IR | Yes | Unpaired t-test | * | 0.0454 | 14, 13 |
| **Fig 2B (Raptor IHC)** |  | Kruskal-Wallis & Dunn's |  |  |  |
| Raptor-OE vs. CCT5-IR | Yes |  | **** | <0.0001 | 17, 13 |
| Raptor-OE vs. Raptor-OE;CCT5-IR | Yes |  | *** | 0.0002 | 17, 10 |
| CCT5-IR vs. Raptor-OE;CCT5-IR | No |  | ns | >0.9999 | 15, 10 |
| WT vs. CCT5-IR | Yes |  | * | 0.0355 | 15, 13 |
| WT vs. Raptor-OE;CCT5-IR | Yes |  | ns | 0.2341 | 15, 10 |
| **Fig 2C (P-S6k IHC)** |  | Kruskal-Wallis & Dunn's |  |  |  |
| WT vs. CCT5-IR | No |  | **** | <0.0001 | 13, 10 |
| WT vs. S6k-OE;CCT5-IR | Yes |  | **** | <0.0001 | 13, 11 |
| CCT5-IR vs. S6k-OE;CCT5-IR | No |  | ns | 0.9773 | 10, 11 |
| **Fig 2E (TDL)** |  | One-way ANOVA & Šídák's |  |  |  |
| CCT3-IR vs. CCT3-IR;Cul1-IR | Yes |  | ns | 0.9534 | 10, 10 |
| Cullin1-IR vs. S6k-OE;Cul1-IR | Yes |  | ns | 0.7985 | 13, 7 |
| S6k-OE;Cul1-IR vs. S6k-OE | Yes |  | ns | 0.3506 | 7, 10 |
| S6k-OE;CCT5-IR vs. CCT5-IR | Yes |  | ns | 0.0608 | 10, 12 |
| **Fig 3A (IHC acet tub)** |  |  |  |  |  |
| WT vs. CCT3-IR | Yes | One-way ANOVA & Dunnett's | **** | <0.0001 | 14, 11 |
| WT vs. CCT5-IR | Yes | One-way ANOVA & Dunnett's | ** | <0.0001 | 14, 13 |
| WT vs. Raptor-IR | No | One-way ANOVA & Dunnett's | *** | 0.0006 | 14, 9 |
| WT vs. S6k-IR | No | Mann-Whitney | *** | 0.0008 | 14, 12 |
| WT vs. Akt-IR | Yes | One-way ANOVA & Dunnett's | * | 0.0148 | 16, 14 |
| WT vs. Cullin1-IR | Yes | Unpaired t-test | ns | 0.2324 | 14, 13 |
| WT vs. S6k-OE | Yes | One-way ANOVA & Dunnett's | ns | 0.9584 | 10, 9 |
| WT vs. Akt-OE | Yes | One-way ANOVA & Dunnett's | ns | 0.6653 | 16, 12 |
| **Fig 3A (IHC Futsch)** |  |  |  |  |  |
| WT vs. CCT3-IR | Yes | Unpaired t-test | *** | 0.0002 | 13, 12 |
| WT vs. CCT5-IR | Yes | Unpaired t-test | * | 0.0398 | 9, 5 |
| WT vs. Raptor-IR | Yes | Unpaired t-test | ** | 0.0089 | 14, 11 |
| WT vs. S6k-IR | Yes | Unpaired t-test | * | 0.0427 | 13, 13 |
| WT vs. Akt-IR | Yes | One-way ANOVA & Dunnett's | *** | 0.0002 | 14, 13 |
| WT vs. Cullin1-IR | Yes | Unpaired t-test | ns | 0.0767 | 10, 13 |
| WT vs. S6k-OE | Yes | One-way ANOVA & Dunnett's | *** | 0.0004 | 16, 15 |
| WT vs. Akt-OE | Yes | Unpaired t-test | ** | 0.0013 | 10, 13 |
| **Fig 3C (*mCherry::Jupiter*)** |  | Two-Way ANOVA & Tukey's |  |  |  |
| 20 μm: WT vs. CCT3-IR | | | **** | <0.0001 | 12, 10 |
| 20 μm: WT vs. CCT5-IR | | | **** | <0.0001 | 12, 10 |
| 20 μm: WT vs. Raptor-IR | | | **** | <0.0001 | 12, 10 |
| 20 μm: WT vs. S6k-IR | | | ** | 0.0035 | 12, 11 |
| 20 μm: WT vs. Akt-IR | | | ** | 0.0040 | 12, 10 |
| 20 μm: WT vs. Cullin1-IR | | | **** | <0.0001 | 12, 11 |
| 20 μm: WT vs. S6k-OE | | | ns | 0.8037 | 12, 10 |
| 20 μm: WT vs. Akt-OE | | | ns | >0.9999 | 12, 10 |
| 40 μm: WT vs. CCT3-IR | | | **** | <0.0001 | 12, 10 |
| 40 μm: WT vs. CCT5-IR | | | **** | <0.0001 | 12, 10 |
| 40 μm: WT vs. Raptor-IR | | | **** | <0.0001 | 12, 10 |
| 40 μm: WT vs. S6k-IR | | | ** | 0.0035 | 12, 11 |
| 40 μm: WT vs. Akt-IR | | | ** | 0.0040 | 12, 10 |
| 40 μm: WT vs. Cullin1-IR | | | **** | <0.0001 | 12, 11 |
| 40 μm: WT vs. S6k-OE | | | ns | 0.8037 | 12, 10 |
| 40 μm: WT vs. Akt-OE | | | ns | >0.9999 | 12, 10 |
| 60 μm: WT vs. CCT3-IR | | | **** | <0.0001 | 12, 10 |
| 60 μm: WT vs. CCT5-IR | | | **** | <0.0001 | 12, 10 |
| 60 μm: WT vs. Raptor-IR | | | **** | <0.0001 | 12, 10 |
| 60 μm: WT vs. S6k-IR | | | ** | 0.0035 | 12, 11 |
| 60 μm: WT vs. Akt-IR | | | ** | 0.0040 | 12, 10 |
| 60 μm: WT vs. Cullin1-IR | | | **** | <0.0001 | 12, 11 |
| 60 μm: WT vs. S6k-OE | | | ns | 0.8037 | 12, 10 |
| 60 μm: WT vs. Akt-OE | | | ns | >0.9999 | 12, 10 |
| 80 μm: WT vs. CCT3-IR | | | **** | <0.0001 | 12, 10 |
| 80 μm: WT vs. CCT5-IR | | | **** | <0.0001 | 12, 10 |
| 80 μm: WT vs. Raptor-IR | | | **** | <0.0001 | 12, 10 |
| 80 μm: WT vs. S6k-IR | | | ** | 0.0035 | 12, 11 |
| 80 μm: WT vs. Akt-IR | | | ** | 0.0040 | 12, 10 |
| 80 μm: WT vs. Cullin1-IR | | | **** | <0.0001 | 12, 11 |
| 80 μm: WT vs. S6k-OE | | | ns | 0.8037 | 12, 10 |
| 80 μm: WT vs. Akt-OE | | | ns | >0.9999 | 12, 10 |
| 100 μm: WT vs. CCT3-IR | | | **** | <0.0001 | 12, 10 |
| 100 μm: WT vs. CCT5-IR | | | **** | <0.0001 | 12, 10 |
| 100 μm: WT vs. Raptor-IR | | | **** | <0.0001 | 12, 10 |
| 100 μm: WT vs. S6k-IR | | | ** | 0.0035 | 12, 11 |
| 100 μm: WT vs. Akt-IR | | | ** | 0.0040 | 12, 10 |
| 100 μm: WT vs. Cullin1-IR | | | **** | <0.0001 | 12, 11 |
| 100 μm: WT vs. S6k-OE | | | ns | 0.8037 | 12, 10 |
| 100 μm: WT vs. Akt-OE | | | ns | >0.9999 | 12, 10 |
| 120 μm: WT vs. CCT3-IR | | | **** | <0.0001 | 12, 10 |
| 120 μm: WT vs. CCT5-IR | | | **** | <0.0001 | 12, 10 |
| 120 μm: WT vs. Raptor-IR | | | **** | <0.0001 | 12, 10 |
| 120 μm: WT vs. S6k-IR | | | ** | 0.0035 | 12, 11 |
| 120 μm: WT vs. Akt-IR | | | ** | 0.0040 | 12, 10 |
| 120 μm: WT vs. Cullin1-IR | | | **** | <0.0001 | 12, 11 |
| 120 μm: WT vs. S6k-OE | | | ns | 0.8037 | 12, 10 |
| 120 μm: WT vs. Akt-OE | | | ns | >0.9999 | 12, 10 |
| 140 μm: WT vs. CCT3-IR | | | **** | <0.0001 | 12, 10 |
| 140 μm: WT vs. CCT5-IR | | | **** | <0.0001 | 12, 10 |
| 140 μm: WT vs. Raptor-IR | | | **** | <0.0001 | 12, 10 |
| 140 μm: WT vs. S6k-IR | | | ** | 0.0035 | 12, 11 |
| 140 μm: WT vs. Akt-IR | | | ** | 0.0040 | 12, 10 |
| 140 μm: WT vs. Cullin1-IR | | | **** | <0.0001 | 12, 11 |
| 140 μm: WT vs. S6k-OE | | | ns | 0.8037 | 12, 10 |
| 140 μm: WT vs. Akt-OE | | | ns | >0.9999 | 12, 10 |
| 160 μm: WT vs. CCT3-IR | | | **** | <0.0001 | 12, 10 |
| 160 μm: WT vs. CCT5-IR | | | **** | <0.0001 | 12, 10 |
| 160 μm: WT vs. Raptor-IR | | | **** | <0.0001 | 12, 10 |
| 160 μm: WT vs. S6k-IR | | | ** | 0.0035 | 12, 11 |
| 160 μm: WT vs. Akt-IR | | | ** | 0.0040 | 12, 10 |
| 160 μm: WT vs. Cullin1-IR | | | **** | <0.0001 | 12, 11 |
| 160 μm: WT vs. S6k-OE | | | ns | 0.8037 | 12, 10 |
| 160 μm: WT vs. Akt-OE | | | ns | >0.9999 | 12, 10 |
| 180 μm: WT vs. CCT3-IR | | | **** | <0.0001 | 12, 10 |
| 180 μm: WT vs. CCT5-IR | | | **** | <0.0001 | 12, 10 |
| 180 μm: WT vs. Raptor-IR | | | **** | <0.0001 | 12, 10 |
| 180 μm: WT vs. S6k-IR | | | ** | 0.0035 | 12, 11 |
| 180 μm: WT vs. Akt-IR | | | ** | 0.0040 | 12, 10 |
| 180 μm: WT vs. Cullin1-IR | | | **** | <0.0001 | 12, 11 |
| 180 μm: WT vs. S6k-OE | | | ns | 0.8037 | 12, 10 |
| 180 μm: WT vs. Akt-OE | | | ns | >0.9999 | 12, 10 |
| 200 μm: WT vs. CCT3-IR | | | **** | <0.0001 | 12, 10 |
| 200 μm: WT vs. CCT5-IR | | | **** | <0.0001 | 12, 10 |
| 200 μm: WT vs. Raptor-IR | | | **** | <0.0001 | 12, 10 |
| 200 μm: WT vs. S6k-IR | | | ** | 0.0035 | 12, 11 |
| 200 μm: WT vs. Akt-IR | | | ** | 0.0040 | 12, 10 |
| 200 μm: WT vs. Cullin1-IR | | | **** | <0.0001 | 12, 11 |
| 200 μm: WT vs. S6k-OE | | | ns | 0.8037 | 12, 10 |
| 200 μm: WT vs. Akt-OE | | | ns | >0.9999 | 12, 10 |
| 220 μm: WT vs. CCT3-IR | | | **** | <0.0001 | 12, 10 |
| 220 μm: WT vs. CCT5-IR | | | **** | <0.0001 | 12, 10 |
| 220 μm: WT vs. Raptor-IR | | | **** | <0.0001 | 12, 10 |
| 220 μm: WT vs. S6k-IR | | | ** | 0.0035 | 12, 11 |
| 220 μm: WT vs. Akt-IR | | | ** | 0.0040 | 12, 10 |
| 220 μm: WT vs. Cullin1-IR | | | **** | <0.0001 | 12, 11 |
| 220 μm: WT vs. S6k-OE | | | ns | 0.8037 | 12, 10 |
| 220 μm: WT vs. Akt-OE | | | ns | >0.9999 | 12, 10 |
| 240 μm: WT vs. CCT3-IR | | | **** | <0.0001 | 12, 10 |
| 240 μm: WT vs. CCT5-IR | | | **** | <0.0001 | 12, 10 |
| 240 μm: WT vs. Raptor-IR | | | **** | <0.0001 | 12, 10 |
| 240 μm: WT vs. S6k-IR | | | ** | 0.0035 | 12, 11 |
| 240 μm: WT vs. Akt-IR | | | ** | 0.0040 | 12, 10 |
| 240 μm: WT vs. Cullin1-IR | | | **** | <0.0001 | 12, 11 |
| 240 μm: WT vs. S6k-OE | | | ns | 0.8037 | 12, 10 |
| 240 μm: WT vs. Akt-OE | | | ns | >0.9999 | 12, 10 |
| 260 μm: WT vs. CCT3-IR | | | **** | <0.0001 | 12, 10 |
| 260 μm: WT vs. CCT5-IR | | | **** | <0.0001 | 12, 10 |
| 260 μm: WT vs. Raptor-IR | | | **** | <0.0001 | 12, 10 |
| 260 μm: WT vs. S6k-IR | | | ** | 0.0035 | 12, 11 |
| 260 μm: WT vs. Akt-IR | | | ** | 0.0040 | 12, 10 |
| 260 μm: WT vs. Cullin1-IR | | | **** | <0.0001 | 12, 11 |
| 260 μm: WT vs. S6k-OE | | | ns | 0.8037 | 12, 10 |
| 260 μm: WT vs. Akt-OE | | | ns | >0.9999 | 12, 10 |
| 280 μm: WT vs. CCT3-IR | | | **** | <0.0001 | 12, 10 |
| 280 μm: WT vs. CCT5-IR | | | **** | <0.0001 | 12, 10 |
| 280 μm: WT vs. Raptor-IR | | | **** | <0.0001 | 12, 10 |
| 280 μm: WT vs. S6k-IR | | | ** | 0.0035 | 12, 11 |
| 280 μm: WT vs. Akt-IR | | | ** | 0.0040 | 12, 10 |
| 280 μm: WT vs. Cullin1-IR | | | **** | <0.0001 | 12, 11 |
| 280 μm: WT vs. S6k-OE | | | ns | 0.8037 | 12, 10 |
| 280 μm: WT vs. Akt-OE | | | ns | >0.9999 | 12, 10 |
| 300 μm: WT vs. CCT3-IR | | | **** | <0.0001 | 12, 10 |
| 300 μm: WT vs. CCT5-IR | | | **** | <0.0001 | 12, 10 |
| 300 μm: WT vs. Raptor-IR | | | **** | <0.0001 | 12, 10 |
| 300 μm: WT vs. S6k-IR | | | ** | 0.0035 | 12, 11 |
| 300 μm: WT vs. Akt-IR | | | ** | 0.0040 | 12, 10 |
| 300 μm: WT vs. Cullin1-IR | | | **** | <0.0001 | 12, 11 |
| 300 μm: WT vs. S6k-OE | | | ns | 0.8037 | 12, 10 |
| 300 μm: WT vs. Akt-OE | | | ns | >0.9999 | 12, 10 |
| 320 μm: WT vs. CCT3-IR | | | **** | <0.0001 | 12, 10 |
| 320 μm: WT vs. CCT5-IR | | | **** | <0.0001 | 12, 10 |
| 320 μm: WT vs. Raptor-IR | | | **** | <0.0001 | 12, 10 |
| 320 μm: WT vs. S6k-IR | | | ** | 0.0035 | 12, 11 |
| 320 μm: WT vs. Akt-IR | | | ** | 0.0040 | 12, 10 |
| 320 μm: WT vs. Cullin1-IR | | | **** | <0.0001 | 12, 11 |
| 320 μm: WT vs. S6k-OE | | | ns | 0.8037 | 12, 10 |
| 320 μm: WT vs. Akt-OE | | | ns | >0.9999 | 12, 10 |
| 340 μm: WT vs. CCT3-IR | | | **** | <0.0001 | 12, 10 |
| 340 μm: WT vs. CCT5-IR | | | **** | <0.0001 | 12, 10 |
| 340 μm: WT vs. Raptor-IR | | | **** | <0.0001 | 12, 10 |
| 340 μm: WT vs. S6k-IR | | | ** | 0.0035 | 12, 11 |
| 340 μm: WT vs. Akt-IR | | | ** | 0.0040 | 12, 10 |
| 340 μm: WT vs. Cullin1-IR | | | **** | <0.0001 | 12, 11 |
| 340 μm: WT vs. S6k-OE | | | ns | 0.8037 | 12, 10 |
| 340 μm: WT vs. Akt-OE | | | ns | >0.9999 | 12, 10 |
| 360 μm: WT vs. CCT3-IR | | | **** | <0.0001 | 12, 10 |
| 360 μm: WT vs. CCT5-IR | | | **** | <0.0001 | 12, 10 |
| 360 μm: WT vs. Raptor-IR | | | **** | <0.0001 | 12, 10 |
| 360 μm: WT vs. S6k-IR | | | ** | 0.0035 | 12, 11 |
| 360 μm: WT vs. Akt-IR | | | ** | 0.0040 | 12, 10 |
| 360 μm: WT vs. Cullin1-IR | | | **** | <0.0001 | 12, 11 |
| 360 μm: WT vs. S6k-OE | | | ns | 0.8037 | 12, 10 |
| 360 μm: WT vs. Akt-OE | | | ns | >0.9999 | 12, 10 |
| 380 μm: WT vs. CCT3-IR | | | **** | <0.0001 | 12, 10 |
| 380 μm: WT vs. CCT5-IR | | | **** | <0.0001 | 12, 10 |
| 380 μm: WT vs. Raptor-IR | | | **** | <0.0001 | 12, 10 |
| 380 μm: WT vs. S6k-IR | | | ** | 0.0035 | 12, 11 |
| 380 μm: WT vs. Akt-IR | | | ** | 0.0040 | 12, 10 |
| 380 μm: WT vs. Cullin1-IR | | | **** | <0.0001 | 12, 11 |
| 380 μm: WT vs. S6k-OE | | | ns | 0.8037 | 12, 10 |
| 380 μm: WT vs. Akt-OE | | | ns | >0.9999 | 12, 10 |
| 400 μm: WT vs. CCT3-IR | | | **** | <0.0001 | 12, 10 |
| 400 μm: WT vs. CCT5-IR | | | **** | <0.0001 | 12, 10 |
| 400 μm: WT vs. Raptor-IR | | | **** | <0.0001 | 12, 10 |
| 400 μm: WT vs. S6k-IR | | | ** | 0.0035 | 12, 11 |
| 400 μm: WT vs. Akt-IR | | | ** | 0.0040 | 12, 10 |
| 400 μm: WT vs. Cullin1-IR | | | **** | <0.0001 | 12, 11 |
| 400 μm: WT vs. S6k-OE | | | ns | 0.8037 | 12, 10 |
| 400 μm: WT vs. Akt-OE | | | ns | >0.9999 | 12, 10 |
| 420 μm: WT vs. CCT3-IR | | | **** | <0.0001 | 12, 10 |
| 420 μm: WT vs. CCT5-IR | | | **** | <0.0001 | 12, 10 |
| 420 μm: WT vs. Raptor-IR | | | **** | <0.0001 | 12, 10 |
| 420 μm: WT vs. S6k-IR | | | ** | 0.0035 | 12, 11 |
| 420 μm: WT vs. Akt-IR | | | ** | 0.0040 | 12, 10 |
| 420 μm: WT vs. Cullin1-IR | | | **** | <0.0001 | 12, 11 |
| 420 μm: WT vs. S6k-OE | | | ns | 0.8037 | 12, 10 |
| 420 μm: WT vs. Akt-OE | | | ns | >0.9999 | 12, 10 |
| 440 μm: WT vs. CCT3-IR | | | **** | <0.0001 | 12, 10 |
| 440 μm: WT vs. CCT5-IR | | | **** | <0.0001 | 12, 10 |
| 440 μm: WT vs. Raptor-IR | | | **** | <0.0001 | 12, 10 |
| 440 μm: WT vs. S6k-IR | | | ** | 0.0035 | 12, 11 |
| 440 μm: WT vs. Akt-IR | | | ** | 0.0040 | 12, 10 |
| 440 μm: WT vs. Cullin1-IR | | | **** | <0.0001 | 12, 11 |
| 440 μm: WT vs. S6k-OE | | | ns | 0.8037 | 12, 10 |
| 440 μm: WT vs. Akt-OE | | | ns | >0.9999 | 12, 10 |
| 460 μm: WT vs. CCT3-IR | | | **** | <0.0001 | 12, 10 |
| 460 μm: WT vs. Raptor-IR | | | **** | <0.0001 | 12, 10 |
| 460 μm: WT vs. Akt-IR | | | ** | 0.0040 | 12, 10 |
| 460 μm: WT vs. Cullin1-IR | | | **** | <0.0001 | 12, 11 |
| 460 μm: WT vs. S6k-OE | | | ns | 0.8037 | 12, 11 |
| 460 μm: WT vs. Akt-OE | | | ns | >0.9999 | 12, 10 |
| 480 μm: WT vs. Akt-IR | | | ** | 0.0040 | 12, 10 |
| 480 μm: WT vs. Cullin1-IR | | | **** | <0.0001 | 12, 11 |
| 480 μm: WT vs. S6k-OE | | | ns | 0.8037 | 12, 11 |
| 480 μm: WT vs. Akt-OE | | | ns | >0.9999 | 12, 10 |
| 500 μm: WT vs. Akt-IR | | | ** | 0.0040 | 12, 10 |
| 500 μm: WT vs. Cullin1-IR | | | **** | <0.0001 | 12, 11 |
| 500 μm: WT vs. S6k-OE | | | ns | 0.8037 | 12, 11 |
| 500 μm: WT vs. Akt-OE | | | ns | >0.9999 | 12, 10 |
| **Fig 4D (HTT *mCherry::Jupiter*)** |  | Two-Way ANOVA & Tukey's |  |  |  |
| 40 μm: HTT20 vs. HTT50 | | | ns | 0.9996 | 10,9 |
| 40 μm: HTT20 vs. HTT93 | | | **** | <0.0001 | 10,11 |
| 80 μm: HTT20 vs. HTT50 | | | ns | 0.9996 | 10,9 |
| 80 μm: HTT20 vs. HTT93 | | | **** | <0.0001 | 10,11 |
| 120 μm: HTT20 vs. HTT50 | | | ns | 0.9996 | 10,9 |
| 120 μm: HTT20 vs. HTT93 | | | **** | <0.0001 | 10,11 |
| 160 μm: HTT20 vs. HTT50 | | | ns | 0.9996 | 10,9 |
| 160 μm: HTT20 vs. HTT93 | | | **** | <0.0001 | 10,11 |
| 200 μm: HTT20 vs. HTT50 | | | ns | 0.9996 | 10,9 |
| 200 μm: HTT20 vs. HTT93 | | | **** | <0.0001 | 10,11 |
| 240 μm: HTT20 vs. HTT50 | | | ns | 0.9996 | 10,9 |
| 240 μm: HTT20 vs. HTT93 | | | **** | <0.0001 | 10,11 |
| 280 μm: HTT20 vs. HTT50 | | | ns | 0.9996 | 10,9 |
| 280 μm: HTT20 vs. HTT93 | | | **** | <0.0001 | 10,11 |
| 320 μm: HTT20 vs. HTT50 | | | ns | 0.9996 | 10,9 |
| 320 μm: HTT20 vs. HTT93 | | | **** | <0.0001 | 10,11 |
| 360 μm: HTT20 vs. HTT50 | | | ns | 0.9996 | 10,9 |
| 360 μm: HTT20 vs. HTT93 | | | **** | <0.0001 | 10,11 |
| 400 μm: HTT20 vs. HTT50 | | | ns | 0.9996 | 10,9 |
| 400 μm: HTT20 vs. HTT93 | | | **** | <0.0001 | 10,11 |
| 440 μm: HTT20 vs. HTT50 | | | ns | 0.9996 | 10,9 |
| 440 μm: HTT20 vs. HTT93 | | | **** | <0.0001 | 10,11 |
| **Fig 4E (Pearson's)** |  | One-way ANOVA & Šídák's |  |  |  |
| WT (HA) GAL4 "off" vs. HTT120 (HA) GAL4 "off" | Yes |  | ns | 0.9965 | 11, 9 |
| WT control (HA) GAL4 “off” vs. HTT120 WT HTT GAL4 “off” | Yes |  | ns | 0.4664 | 11, 10 |
| HTT120 (HA) GAL4 “off” vs. HTT120 (HA) GAL4 “on” | Yes |  | ** | 0.0045 | 9, 11 |
| HTT120 WT HTT GAL4 “off” vs. HTT120 (HA) GAL4 “on” | Yes |  | ns | 0.5580 | 10, 11 |
| **Fig 4F (Manders Soma)** |  |  |  |  |  |
| WT HTT v HTT120HA 48 hr | No | Mann-Whitney | *** | 0.0007 | 10, 11 |
| **Fig 4G (HTT TDL)** |  | One-way ANOVA & Šídák’s |  |  |  |
| WT vs. HTTQ25 | Yes |  | ** | 0.0042 | 11, 11 |
| WT vs. HTT96 | Yes |  | * | 0.0211 | 11, 16 |
| WT vs. HTTQ96;CCT5-IR | Yes |  | **** | <0.0001 | 11, 16 |
| WT vs. HTTQ96;Cul1-IR | Yes |  | ns | 0.8400 | 11, 9 |
| WT vs. HTT25;CCT5-IR | Yes |  | **** | <0.0001 | 11, 9 |
| WT vs. HTTQ25;Cul1-IR | Yes |  | * | 0.0383 | 11, 9 |
| HTTQ25 vs. HTTQ25;CCT5-IR | Yes |  | **** | <0.0001 | 11, 9 |
| HTTQ25 vs. HTTQ25;Cul1-IR | Yes |  | ns | >0.9999 | 11, 9 |
| HTTQ96 vs. HTTQ96;CCT5-IR | Yes |  | *** | 0.0004 | 16, 16 |
| HTTQ96 vs. HTTQ96;Cul1-IR | Yes |  | *** | 0.0002 | 16, 9 |
| HTTQ96;CCT5-IR vs. HTTQ25;CCT5-IR | Yes |  | ns | 0.9993 | 16, 9 |
| HTTQ96;Cul1-IR vs. HTTQ25;Cul1-IR | Yes |  | ns | 0.8662 | 9, 9 |
| **Fig S1A (CCT TDL)** |  | One-way ANOVA & Dunnett’s |  |  |  |
| WT vs. CCT1-IR | Yes |  | **** | <0.0001 | 13, 10 |
| WT vs. CCT2-IR | Yes |  | ns | 0.1586 | 13, 10 |
| WT vs. CCT4-IR | Yes |  | ns | 0.6527 | 13, 10 |
| WT vs. CCT6-IR | Yes |  | **** | <0.0001 | 13, 10 |
| WT vs. CCT7-IR | Yes |  | * | 0.0388 | 13, 10 |
| WT vs. CCT8-IR | Yes |  | **** | <0.0001 | 13, 11 |
| **Fig S1B (CCT4 TDL)** |  |  |  |  |  |
| 40A FRT vs. CCT4<sup>KG09280</sup> | No | Mann-Whitney Test | *** | 0.0003 | 11, 8 |
| **Fig S1D (CCT Sholl Max)** |  | One-way ANOVA & Dunnett’s |  |  |  |
| WT vs. CCT1-IR | Yes |  | ** | 0.0025 | 9, 10 |
| WT vs. CCT2-IR | Yes |  | ns | 0.9665 | 9, 10 |
| WT vs. CCT4-IR | Yes |  | ns | 0.5974 | 9, 11 |
| WT vs. CCT6-IR | Yes |  | * | 0.0153 | 9, 10 |
| WT vs. CCT7-IR | Yes |  | ns | 0.3998 | 9, 10 |
| WT vs. CCT8-IR | Yes |  | * | 0.0471 | 9, 11 |
| **Fig S1D (CCT Sholl Radius)** |  | One-way ANOVA & Dunnett's |  |  |  |
| WT vs. CCT1-IR | Yes |  | *** | 0.0001 | 9, 10 |
| WT vs. CCT2-IR | Yes |  | ns | 0.2714 | 9, 10 |
| WT vs. CCT4-IR | Yes |  | ns | 0.3769 | 9, 11 |
| WT vs. CCT6-IR | Yes |  | *** | 0.0007 | 9, 10 |
| WT vs. CCT7-IR | Yes |  | ns | >0.9999 | 9, 10 |
| WT vs. CCT8-IR | Yes |  | ns | 0.1483 | 9, 11 |
| **Fig S1E (CCT5 IHC)** |  | One-way ANOVA & Šídák's |  |  |  |
| WT vs. CCT4-IR | Yes |  | **** | <0.0001 | 29, 23 |
| WT vs. CCT5-IR | Yes |  | *** | 0.0002 | 29, 16 |
| WT vs. CCT4-IR;CCT5-IR | Yes |  | **** | <0.0001 | 29, 18 |
| CCT4-IR vs. CCT4-IR;CCT5-IR | Yes |  | ** | 0.0021 | 23, 18 |
| CCT5-IR vs. CCT4-IR;CCT5-IR | Yes |  | *** | 0.0004 | 16, 18 |
| **Fig S1F (AEL TDL)** |  | One-way ANOVA & Šídák's |  |  |  |
| 24 hr: mean of genetic controls vs. CCT3-IR | Yes |  | ns | 0.4890 | 17, 15 |
| 24 hr: mean of genetic controls vs. CCT5-IR | Yes |  | ns | 0.4424 | 12, 13 |
| 48 hr: mean of genetic controls vs. CCT3-IR | Yes |  | ns | 0.1891 | 18, 12 |
| 48 hr: mean of genetic controls vs. CCT5-IR | Yes |  | ns | 0.6169 | 10, 12 |
| 72 hr: mean of genetic controls vs. CCT3-IR | Yes |  | *** | 0.0004 | 11, 14 |
| 72 hr: mean of genetic controls vs. CCT5-IR | Yes |  | * | 0.0104 | 10, 12 |
| 96 hr: mean of genetic controls vs. CCT3-IR | Yes |  | **** | <0.0001 | 10, 10 |
| 96 hr: mean of genetic controls vs. CCT5-IR | Yes |  | **** | <0.0001 | 12, 12 |
| 72 hr CCT3-IR vs. 96 hr CCT3-IR | Yes |  | ns | 0.6523 | 14, 10 |
| 72 hr CCT5-IR vs. 96 hr CCT5-IR | Yes |  | ns | 0.5468 | 12, 12 |
| **Fig S1G (CCT OE TDL)** |  | One-way ANOVA & Šídák's |  |  |  |
| WT vs. CCT2-OE | Yes |  | ns | 0.9917 | 13, 10 |
| WT vs. drosCCT4 | Yes |  | ns | 0.6067 | 13, 10 |
| dcas9 vs. CCT5 TOE | Yes |  | ns | 0.2882 | 13, 11 |
| **Fig S2A (Raptor fluorescence IHC)** |  |  |  |  |  |
| WT vs. CCT5-IR | Yes | One-way ANOVA & Dunnett's | **** | <0.0001 | 11, 14 |
| WT vs. Raptor-IR | Yes | One-way ANOVA & Dunnett's | **** | <0.0001 | 16, 16 |
| WT vs. Raptor-OE | Yes | One-way ANOVA & Dunnett's | *** | 0.0005 | 16, 14 |
| **Fig S2B (S6k IHC)** |  | Unpaired t-test |  |  |  |
| WT vs. S6k-IR | Yes |  | ** | 0.0071 | 9, 8 |
| **Fig S2C (P-Akt IHC)** |  | Unpaired t-test |  |  |  |
| WT vs. Akt-IR | Yes |  | **** | <0.0001 | 11, 12 |
| **Fig S2D (Cullin1 IHC)** |  | Unpaired t-test |  |  |  |
| WT vs. Cullin1-IR | Yes |  | ** | 0.0013 | 15, 14 |
| **Fig S2E (BtubIIA fluorescence)** |  |  |  |  |  |
| WT vs. CCT3-IR | Yes | One-way ANOVA & Dunnett's | **** | <0.0001 | 14, 11 |
| WT vs. CCT5-IR | Yes | One-way ANOVA & Dunnett's | **** | <0.0001 | 14, 15 |
| WT vs. S6k-IR | Yes | Unpaired t-test | * | 0.0339 | 13, 13 |
| WT vs. Raptor-IR | Yes | Unpaired t-test | ** | 0.0078 | 14, 11 |
| WT vs. Akt-IR | Yes | One-way ANOVA & Dunnett's | ** | 0.0055 | 16, 14 |
| WT vs. Akt-OE | Yes | One-way ANOVA & Dunnett's | ns | 0.4915 | 16, 12 |
| WT vs. Cullin1-IR | Yes | Unpaired t-test | * | 0.0105 | 10, 13 |
| **Fig S3A (HTT TDL)** |  | One-way ANOVA & Šídák's |  |  |  |
| WT vs. HTTQ20 | Yes |  | ns | >0.9999 | 11, 15 |
| WT vs. HTTQ50 | Yes |  | ns | 0.8793 | 11, 13 |
| WT vs. HTTQ93 | Yes |  | **** | <0.0001 | 11, 18 |
| WT vs. HTTQ120 | Yes |  | **** | <0.0001 | 11, 15 |
| **Fig S3C (HTT Futsch)** |  | One-way ANOVA & Tukey's |  |  |  |
| WT vs. HTT96 | Yes |  | **** | <0.0001 | 11, 11 |
| WT vs. HTT96;CCT5-IR | Yes |  | **** | <0.0001 | 11, 9 |
| HTT96 vs. HTT96;CCT5-IR | Yes |  | ns | >0.9999 | 11, 9 |
| **Fig S3D (WT HTT)** |  |  |  |  |  |
| WT vs. CCT5-IR | Yes | Unpaired t-test | *** | 0.0002 | 11, 10 |
| **Fig S3F (M2 Dendrites)** |  |  |  |  |  |
| WT HTT v HTT120HA 48 hr | Yes | Unpaired t-test | **** | <0.0001 | 11, 9 |
| **Fig S3G (WT HTT IHC)** |  | One-way ANOVA & Tukey's |  |  |  |
| WT vs. HTT25-Cer | Yes |  | * | 0.0117 | 11, 11 |
| WT vs. HTT96-Cer | Yes |  | **** | <0.0001 | 11, 11 |
| **Fig S3H (# mHTT IB)** |  | Kruskal-Wallis & Dunn's |  |  |  |
| HTT96 vs. HTT96;CCT5-IR | No |  | ns | >0.9999 | 15, 14 |
| HTT96 vs. HTT96;Cul1-IR | No |  | ns | 0.7909 | 15, 14 |
| **Fig S3I (area mHTT IB)** |  | Kruskal-Wallis & Dunn's |  |  |  |
| HTT96 vs. HTT96;CCT5-IR | No |  | ns | 0.1699 | 16, 16 |
| HTT96 vs. HTT96;Cul1-IR | No |  | ns | >0.9999 | 16, 15 |

## Notes

**CONFLICT OF INTEREST** The authors declare no competing financial interests.

### Competing Interest Statement

The authors have declared no competing interest.

## REFERENCES

Arshadi C, Günther U, Eddison M, Harrington KIS, Ferreira TA (2021) SNT: a unifying toolbox for quantification of neuronal anatomy. Nat Methods 18:374–377.

Barnat M, Le Friec J, Benstaali C, Humbert S (2017) Huntingtin-Mediated Multipolar-Bipolar Transition of Newborn Cortical Neurons Is Critical for Their Postnatal Neuronal Morphology. Neuron 93:99– 114.

Berger Z, Ravikumar B, Menzies FM, Garcia Oroz L, Underwood BR, Pangalos MN, Schmitt I, Wullner U, Evert BO, O’kane CJ, Rubinsztein DC (2006) Rapamycin alleviates toxicity of different aggregate-prone proteins. Human Molecular Genetics 15:433–442.

Bertrand M, Decoville M, Meudal H, Birman S, Landon C (2020) Metabolomic nuclear magnetic resonance studies at presymptomatic and symptomatic stages of huntington’s disease on a drosophila model. Journal of Proteome Research 19:4034–4045.

Bhattacharjee S, Lottes EN, Nanda S, Golshir A, Patel AA, Ascoli GA, Cox DN (2022) PP2A phosphatase regulates cell-type specific cytoskeletal organization to drive dendrite diversity. Front Mol Neurosci 15:926567.

Bhutani S, Das A, Maheshwari M, Lakhotia SC, Jana NR (2012) Dysregulation of core components of SCF complex in poly-glutamine disorders. Cell Death Dis 3:e428–e428.

Bolte S, Cordelières FP (2006) A guided tour into subcellular colocalization analysis in light microscopy. Journal of Microscopy 224:213–232.

Brackley KI, Grantham J (2009) Activities of the chaperonin containing TCP-1 (CCT): Implications for cell cycle progression and cytoskeletal organisation. Cell Stress and Chaperones 14:23–31.

Burillo J, Marqués P, Jiménez B, González-Blanco C, Benito M, Guillén C (2021) Insulin Resistance and Diabetes Mellitus in Alzheimer’s Disease. Cells 10:1–42.

Chen X-Q, Fang F, Florio JB, Rockenstein E, Masliah E, Mobley WC, Rissman RA, Wu C (2018) T-complex protein 1-ring complex enhances retrograde axonal transport by modulating tau phosphorylation. Traffic:1–14.

Chen ZS, Wong AKY, Cheng TC, Koon AC, Chan HYE (2019) FipoQ/FBXO33, a Cullin-1-based ubiquitin ligase complex component modulates ubiquitination and solubility of polyglutamine disease protein. Journal of Neurochemistry 149:781–798.

Clark SG, Graybeal LL, Bhattacharjee S, Thomas C, Bhattacharya S, Cox DN (2018) Basal autophagy is required for promoting dendritic terminal branching in drosophila sensory neurons. PLoS ONE 13:e0206743.

Cordelières FP, Bolte S (2014) Chapter 21 - Experimenters’ guide to colocalization studies: Finding a way through indicators and quantifiers, in practice. In: Methods in Cell Biology (Waters JC, Wittman T, eds), pp 395–408 Quantitative Imaging in Cell Biology. Academic Press.

Craft S (2009) The Role of Metabolic Disorders in Alzheimer Disease and Vascular Dementia: Two Roads Converged. Archives of Neurology 66:300–305.

Cuéllar J, Ludlam WG, Tensmeyer NC, Aoba T, Dhavale M, Santiago C, Bueno-Carrasco MT, Mann MJ, Plimpton RL, Makaju A, Franklin S, Willardson BM, Valpuesta JM (2019) Structural and functional analysis of the role of the chaperonin CCT in mTOR complex assembly. Nature Communications 10:1–14.

Das R, Bhattacharjee S, Patel AA, Harris JM, Bhattacharya S, Letcher JM, Clark SG, Nanda S, Iyer EPR, Ascoli GA, Cox DN (2017) Dendritic cytoskeletal architecture is modulated by combinatorial transcriptional regulation in Drosophila melanogaster. Genetics 207:1401–1421.

Dewey EH (1900) The no-breakfast plan and the fasting-cure, Fourth Edition. New York city: The Health culture co.

Diaz-Ruiz O, Zapata A, Shan L, Zhang Y, Tomac AC, Malik N, Cruz F de la, Bäckman CM (2009) Selective Deletion of PTEN in Dopamine Neurons Leads to Trophic Effects and Adaptation of Striatal Medium Spiny Projecting Neurons. PLOS ONE 4:e7027.

Eshun-Wilson L, Zhang R, Portran D, Nachury MV, Toso DB, Löhr T, Vendruscolo M, Bonomi M, Fraser JS, Nogales E (2019) Effects of α-tubulin acetylation on microtubule structure and stability. Proc Natl Acad Sci U S A 116:10366–10371.

Feng L, Zhao T, Kim J (2015) neuTube 1.0: A New Design for Efficient Neuron Reconstruction Software Based on the SWC Format. eNeuro 2.

Ferreira TA, Blackman AV, Oyrer J, Jayabal S, Chung AJ, Watt AJ, Sjöström PJ, van Meyel DJ (2014) Neuronal morphometry directly from bitmap images. Nat Methods 11:982–984.

Fingar DC, Salama S, Tsou C, Harlow E, Blenis J (2002) Mammalian cell size is controlled by mTOR and its downstream targets S6K1 and 4EBP1/eIF4E. Genes & Development 16:1472–1487.

Freund A, Zhong FL, Venteicher AS, Meng Z, Veenstra TD, Frydman J, Artandi SE (2014) Proteostatic control of telomerase function through TRiC-mediated folding of TCAB1. Cell 159:1389–1403.

Gerez JA, Prymaczok NC, Rockenstein E, Herrmann US, Schwarz P, Adame A, Enchev RI, Courtheoux T, Boersema PJ, Riek R, Peter M, Aguzzi A, Masliah E, Picotti P (2019) A cullin-RING ubiquitin ligase targets exogenous α-synuclein and inhibits Lewy body–like pathology. Science Translational Medicine 11:eaau6722.

Gestaut D, Zhao Y, Park J, Ma B, Leitner A, Collier M, Pintilie G, Roh S-H, Chiu W, Frydman J (2022) Structural visualization of the tubulin folding pathway directed by human chaperonin TRiC/CCT. Cell 185:4770–4787.e20.

Grantham J, Brackley KI, Willison KR (2006) Substantial CCT activity is required for cell cycle progression and cytoskeletal organization in mammalian cells. Experimental Cell Research 312:2309–2324.

Grueber WB, Jan LY, Jan YN (2002) Tiling of the Drosophila epidermis by multidendritic sensory neurons. Development 129:2867–2878.

Hać A, Pierzynowska K, Herman-Antosiewicz A (2021) S6k1 is indispensible for stress-induced microtubule acetylation and autophagic flux. Cells 10.

Hummel T, Krukkert K, Roos J, Davis G, Klämbt C (2000) Drosophila Futsch/22C10 Is a MAP1B-like Protein Required for Dendritic and Axonal Development. Neuron 26:357–370.

Iyer EPR, Iyer SC, Sullivan L, Wang D, Meduri R, Graybeal LL, Cox DN (2013a) Functional Genomic Analyses of Two Morphologically Distinct Classes of Drosophila Sensory Neurons: Post-Mitotic Roles of Transcription Factors in Dendritic Patterning. PLoS ONE 8:e72434.

Iyer SC, Iyer EPR, Meduri R, Rubaharan M, Kuntimaddi A, Karamsetty M, Cox DN (2013b) Cut, via CrebA, transcriptionally regulates the COPII secretory pathway to direct dendrite development in Drosophila. Journal of Cell Science 126:4732–4745.

Jaworski J, Spangler S, Seeburg DP, Hoogenraad CC, Sheng M (2005) Control of dendritic arborization by the phosphoinositide-3’-kinase-Akt-mammalian target of rapamycin pathway. The Journal of neuroscience : the official journal of the Society for Neuroscience 25:11300–11312.

Jin M, Han W, Liu C, Zang Y, Li J, Wang F, Wang Y, Cong Y (2019) An ensemble of cryo-EM structures of TRiC reveal its conformational landscape and subunit specificity. Proceedings of the National Academy of Sciences of the United States of America 116:19513–19522.

Kanaoka Y, Onodera K, Watanabe K, Hayashi Y, Usui T, Uemura T, Hattori Y (2023) Inter-organ Wingless/Ror/Akt signaling regulates nutrient-dependent hyperarborization of somatosensory neurons. eLife 12:e79461.

Kellar D, Craft S (2020) Brain insulin resistance in Alzheimer’s disease and related disorders: mechanisms and therapeutic approaches. Lancet Neurol 19:758–766.

Khyati, Malik I, Agrawal N, Kumar V (2021) Melatonin and curcumin reestablish disturbed circadian gene expressions and restore locomotion ability and eclosion behavior in Drosophila model of Huntington’s disease. Chronobiology International 38:61–78.

Kim A-R, Choi K-W (2019) TRiC/CCT chaperonins are essential for organ growth by interacting with insulin/TOR signaling in Drosophila. Oncogene 38:4739–4754.

King MA, Hands S, Hafiz F, Mizushima N, Tolkovsky AM, Wyttenbach A (2008) Rapamycin Inhibits Polyglutamine Aggregation Independently of Autophagy by Reducing Protein Synthesis. Mol Pharmacol 73:1052–1063.

Kitamura A, Kubota H, Pack C-G, Matsumoto G, Hirayama S, Takahashi Y, Kimura H, Kinjo M, Morimoto RI, Nagata K (2006) Cytosolic chaperonin prevents polyglutamine toxicity with altering the aggregation state. Nat Cell Biol 8:1163–1169.

Kosillo P, Ahmed KM, Aisenberg EE, Karalis V, Roberts BM, Cragg SJ, Bateup HS (2022) Dopamine neuron morphology and output are differentially controlled by mTORC1 and mTORC2. eLife 11:e75398.

Kosillo P, Doig NM, Ahmed KM, Agopyan-Miu AHCW, Wong CD, Conyers L, Threlfell S, Magill PJ, Bateup HS (2019) Tsc1-mTORC1 signaling controls striatal dopamine release and cognitive flexibility. Nat Commun 10:5426.

Krench M, Littleton JT (2013) Modeling huntington disease in Drosophila: Insights into axonal transport defects and modifiers of toxicity. Fly 7:229–236.

Kumar V, Zhang M-X, Swank MW, Kunz J, Wu G-Y (2005) Regulation of Dendritic Morphogenesis by Ras– PI3K–Akt–mTOR and Ras–MAPK Signaling Pathways. J Neurosci 25:11288–11299.

Lee G, Chung J (2007) Discrete functions of rictor and raptor in cell growth regulation in Drosophila. Biochemical and Biophysical Research Communications 357:1154–1159.

Lee JH, Tecedor L, Chen YH, Mas Monteys A, Sowada MJ, Thompson LM, Davidson BL (2015) Reinstating aberrant mTORC1 activity in Huntington’s disease mice improves disease phenotypes HHS Public Access. Neuron 85:303–315.

Liou AK, Willison KR (1997) Elucidation of the subunit orientation in CCT (chaperonin containing TCP1) from the subunit composition of CCT micro-complexes. EMBO J 16:4311–4316.

Llorca O (2000) Eukaryotic chaperonin CCT stabilizes actin and tubulin folding intermediates in open quasi-native conformations. The EMBO Journal 19:5971–5979.

Mandel SA, Fishman-Jacob T, Youdim MBH (2012a) Genetic reduction of the E3 ubiquitin ligase element, SKP1A and environmental manipulation to emulate cardinal features of Parkinson’s disease. Parkinsonism & Related Disorders 18:S177–S179.

Mandel SA, Fishman-Jacob T, Youdim MBH (2012b) Targeting SKP1, an ubiquitin E3 ligase component found decreased in sporadic Parkinson’s disease. Neurodegener Dis 10:220–223.

McGuire SE, Mao Z, Davis RL (2004) Spatiotemporal gene expression targeting with the TARGET and gene-switch systems in Drosophila. Sci STKE 2004:pl6.

McKinstry SU, Karadeniz YB, Worthington AK, Hayrapetyan VY, Ilcim Ozlu M, Serafin-Molina K, Christopher Risher W, Ustunkaya T, Dragatsis I, Zeitlin S, Yin HH, Eroglu C (2014) Huntingtin is required for normal excitatory synapse development in cortical and striatal circuits. Journal of Neuroscience 34:9455–9472.

Miles WR, Root HF (1922) Psychologic Tests Applied to Diabetic Patients. Archives of Internal Medicine 30:767–777.

Moheet A, Mangia S, Seaquist E (2015) Impact of diabetes on cognitive function and brain structure. Ann N Y Acad Sci 1353:60–71.

Nanda S, Bhattacharjee S, Cox DN, Ascoli GA (2021) An imaging analysis protocol to trace, quantify, and model multi-signal neuron morphology. STAR Protocols 2:100567.

Noormohammadi A, Khodakarami A, Gutierrez-Garcia R, Lee HJ, Koyuncu S, König T, Schindler C, Saez I, Fatima A, Dieterich C, Vilchez D (2016) Somatic increase of CCT8 mimics proteostasis of human pluripotent stem cells and extends C. elegans lifespan. Nature Communications 7:13649.

Pardridge WM, Eisenberg J, Yang J (1985) Human Blood—Brain Barrier Insulin Receptor. Journal of Neurochemistry 44:1771–1778.

Parrish JZ, Xu P, Kim CC, Jan LY, Jan YN (2009) The microRNA bantam functions in epithelial cells to regulate scaling growth of dendrite arbors in Drosophila sensory neurons. Neuron 63:788–802.

Pawson C, Eaton BA, Davis GW (2008) Formin-Dependent Synaptic Growth: Evidence That Dlar Signals via Diaphanous to Modulate Synaptic Actin and Dynamic Pioneer Microtubules. J Neurosci 28:11111–11123.

Pryor WM, Biagioli M, Shahani N, Swarnkar S, Huang WC, Page DT, MacDonald ME, Subramaniam S (2014) Huntingtin promotes mTORC1 signaling in the pathogenesis of Huntington’s disease. Science Signaling 7.

Querfurth H, Lee HK (2021) Mammalian/mechanistic target of rapamycin (mTOR) complexes in neurodegeneration. Molecular Neurodegeneration 16.

Raizada MK (1983) Localization of insulin-like immunoreactivity in the neurons from primary cultures of rat brain. Exp Cell Res 143:351–357.

Ravikumar B, Vacher C, Berger Z, Davies JE, Luo S, Oroz LG, Scaravilli F, Easton DF, Duden R, O’Kane CJ, Rubinsztein DC (2004) Inhibition of mTOR induces autophagy and reduces toxicity of polyglutamine expansions in fly and mouse models of Huntington disease. Nat Genet 36:585– 595.

Roscic A, Baldo B, Crochemore C, Marcellin D, Paganetti P (2011) Induction of autophagy with catalytic mTOR inhibitors reduces huntingtin aggregates in a neuronal cell model. Journal of Neurochemistry 119:398–407.

Sage D, Donati L, Soulez F, Fortun D, Schmit G, Seitz A, Guiet R, Vonesch C, Unser M (2017) DeconvolutionLab2: An open-source software for deconvolution microscopy. Methods 115:28– 41.

Sarkar S, Ravikumar B, Floto RA, Rubinsztein DC (2008) Rapamycin and mTOR-independent autophagy inducers ameliorate toxicity of polyglutamine-expanded huntingtin and related proteinopathies. Cell Death & Differentiation 2009 16:1 16:46–56.

Schulingkamp RJ, Pagano TC, Hung D, Raffa RB (2000) Insulin receptors and insulin action in the brain: review and clinical implications. Neuroscience & Biobehavioral Reviews 24:855–872.

Sergeeva OA, Tran MT, Haase-Pettingell C, King JA (2014) Biochemical characterization of mutants in chaperonin proteins CCT4 and CCT5 associated with hereditary sensory neuropathy. Journal of Biological Chemistry 289:27470–27480.

Shahmoradian SH, Galaz-Montoya JG, Schmid MF, Cong Y, Ma B, Spiess C, Frydman J, Ludtke SJ, Chiu W (2013) TRiC’s tricks inhibit huntingtin aggregation. eLife 2013:710.

Shen K, Calamini B, Fauerbach JA, Ma B, Shahmoradian SH, Serrano Lachapel IL, Chiu W, Lo DC, Frydman J (2016) Control of the structural landscape and neuronal proteotoxicity of mutant Huntingtin by domains flanking the polyQ tract. eLife 5:e18065.

Shimono K, Fujishima K, Nomura T, Ohashi M, Usui T, Kengaku M, Toyoda A, Uemura T (2014) An evolutionarily conserved protein CHORD regulates scaling of dendritic arbors with body size. Scientific Report 4.

Skalecka A, Liszewska E, Bilinski R, Gkogkas C, Khoutorsky A, Malik AR, Sonenberg N, Jaworski J (2016) mTOR Kinase is Needed for the Development and Stabilization of Dendritic Arbors in Newly Born Olfactory Bulb Neurons. Inc Develop Neurobiol 76:1308–1327.

Sontag EM, Joachimiak LA, Tan Z, Tomlinson A, Housman DE, Glabe CG, Potkinj SG, Frydman J, Thompson LM (2013) Exogenous delivery of chaperonin subunit fragment ApiCCT1 modulates mutant Huntingtin cellular phenotypes. Proceedings of the National Academy of Sciences of the United States of America 110:3077–3082.

Subhan I, Siddique YH (2021) Modulation of Huntington’s disease in Drosophila. CNS & Neurological Disorders - Drug Targets 20.

Sulkowski MJ, Iyer SC, Kurosawa MS, Iyer EPR, Cox DN (2011) Turtle Functions Downstream of Cut in Differentially Regulating Class Specific Dendrite Morphogenesis in Drosophila. PLoS One 6:e22611.

Swiech L, Blazejczyk M, Urbanska M, Pietruszka P, Dortland BR, Malik AR, Wulf PS, Hoogenraad CC, Jaworski J (2011) Cellular/Molecular CLIP-170 and IQGAP1 Cooperatively Regulate Dendrite Morphology.

Switon K, Kotulska K, Janusz-Kaminska A, Zmorzynska J, Jaworski J (2017) Molecular neurobiology of mTOR. Neuroscience 341:112–153.

Tam S, Geller R, Spiess C, Frydman J (2006) The chaperonin TRiC controls polyglutamine aggregation and toxicity through subunit-specific interactions. Nature Cell Biology 8:1155–1162.

Tenenbaum CM, Gavis ER (2016) Removal of Drosophila Muscle Tissue from Larval Fillets for Immunofluorescence Analysis of Sensory Neurons and Epidermal Cells. J Vis Exp:54670.

Thomanetz V, Angliker N, Cloëtta D, Lustenberger RM, Schweighauser M, Oliveri F, Suzuki N, Rüegg MA (2013) Ablation of the mTORC2 component rictor in brain or Purkinje cells affects size and neuron morphology. The Journal of Cell Biology 201:293.

Thulasiraman V, Yang C-F, Frydman J (1999) In vivo newly translated polypeptides are sequestered in a protected folding environment.

Trushina E, Heldebrant MP, Perez-Terzic CM, Bortolon R, Kovtun IV, Badger JD, Terzic A, Estévez A, Windebank AJ, Dyer RB, Yao J, McMurray CT (2003) Microtubule destabilization and nuclear entry are sequential steps leading to toxicity in Huntington’s disease. Proc Natl Acad Sci U S A 100:12171–12176.

Urbanska M, Gozdz A, Swiech LJ, Jaworski J (2012) Mammalian Target of Rapamycin Complex 1 (mTORC1) and 2 (mTORC2) Control the Dendritic Arbor Morphology of Hippocampal Neurons. The Journal of Biological Chemistry 287:30240.

Vernizzi L, Paiardi C, Licata G, Vitali T, Santarelli S, Raneli M, Manelli V, Rizzetto M, Gioria M, Pasini ME, Grifoni D, Vanoni MA, Gellera C, Taroni F, Bellosta P (2020) Glutamine Synthetase 1 Increases Autophagy Lysosomal Degradation of Mutant Huntingtin Aggregates in Neurons, Ameliorating Motility in a Drosophila Model for Huntington’s Disease. Cells 9.

Vinayagram A, Kulkarni MM, Sopko R, Sun X, Hu Y, Nand A, Villalta C, Moghimi A, Yang X, Mohr SE, Hong P, Asara JM, Perrimon N (2016) An Integrative Analysis of the InR/PI3K/Akt Network Identifies the Dynamic Response to Insulin Signaling. Cell Reports 16:3062–3074.

Wang YH, Ding ZY, Cheng YJ, Chien CT, Huang ML (2020) An Efficient Screen for Cell-Intrinsic Factors Identifies the Chaperonin CCT and Multiple Conserved Mechanisms as Mediating Dendrite Morphogenesis. Frontiers in Cellular Neuroscience 14:311.

Weiner AT, Lanz MC, Goetschius DJ, Hancock WO, Rolls MM (2016) Kinesin-2 and Apc function at dendrite branch points to resolve microtubule collisions. Cytoskeleton 73:35–44.

Weyhenmeyer JA, Fellows RE (1983) Presence of immunoreactive insulin in neurons cultured from fetal rat brain. Cell Mol Neurobiol 3:81–86.

Williams A, Sarkar S, Cuddon P, Ttofi EK, Saiki S, Siddiqi FH, Jahreiss L, Fleming A, Pask D, Goldsmith P, O’Kane CJ, Floto RA, Rubinsztein DC (2008) Novel targets for Huntington’s disease in an mTOR-independent autophagy pathway. Nat Chem Biol 4:295–305.

Willison KR (2018) The substrate specificity of eukaryotic cytosolic chaperonin CCT. Philosophical Transactions of the Royal Society B: Biological Sciences 373.

Wong JJL, Li S, Lim EKH, Wang Y, Wang C, Zhang H, Kirilly D, Wu C, Liou YC, Wang H, Yu F (2013) A Cullin1-Based SCF E3 Ubiquitin Ligase Targets the InR/PI3K/TOR Pathway to Regulate Neuronal Pruning. PLoS Biology 11.

Wu J, Zhou S-L, Pi L-H, Shi X-J, Ma L-R, Chen Z, Qu M-L, Li X, Nie S-D, Liao D-F, Pei J-J, Wang S (2017) High glucose induces formation of tau hyperphosphorylation via Cav-1-mTOR pathway: A potential molecular mechanism for diabetes-induced cognitive dysfunction. Oncotarget 8:40843–40856.

Zhao X, Chen XQ, Han E, Hu Y, Paik P, Ding Z, Overman J, Lau AL, Shahmoradian SH, Chiu W, Thompson LM, Wu C, Mobley WC (2016) TRiC subunits enhance BDNF axonal transport and rescue striatal atrophy in Huntington’s disease. Proceedings of the National Academy of Sciences of the United States of America 113:E5655–E5664.

